# CNTNAP2 ectodomain, detected in neuronal and CSF sheddomes, modulates Ca^2+^ dynamics and network synchrony

**DOI:** 10.1101/605378

**Authors:** M. Dolores Martin-de-Saavedra, Marc dos Santos, Olga Varea, Benjamin P. Spielman, Ruoqi Gao, Marc Forrest, Kristoffer Myczek, Natalia Khalatyan, Elizabeth A. Hall, Antonio Sanz-Clemente, Davide Comoletti, Stefan F. Lichtenthaler, Jeffrey N. Savas, Peter Penzes

## Abstract

While many neuronal membrane-anchored proteins undergo proteolytic cleavage, little is known about the biological significance of neuronal ectodomain shedding. Using mass spectrometry (MS)-based proteomics, we showed that the neuronal sheddome mirrors human cerebrospinal fluid (hCSF). Among shed synaptic proteins in hCSF was the ectodomain of CNTNAP2 (CNTNAP2-ecto), a risk factor for neurodevelopmental disorders (NDD). Using structured-illumination microscopy (SIM), we mapped the spatial organization of neuronal CNTNAP2-ecto shedding. Using affinity chromatography followed by MS, we identified the ATP2B/PMCA Ca^2+^ extrusion pumps as novel CNTNAP2-ecto binding partners. CNTNAP2-ecto coimmunoprecipitates with PMCA2, a known autism risk factor, and enhances its activity, thereby modulating neuronal Ca^2+^ levels. Finally, we showed that CNTNAP2-ecto regulates neuronal network synchrony in primary cultures and brain slices. These data provide new insights into the biology of synaptic ectodomain shedding and reveal a novel mechanism of regulation of Ca^2+^ homeostasis and neuronal network synchrony.

## INTRODUCTION

Numerous transmembrane proteins undergo proteolytic processing, resulting in the release of a soluble extracellular fragment in a process called ectodomain shedding. Ectodomain shedding plays a widespread role in development and disease, including in the central nervous system (Lichtenthaler et al., 2018). Conversely, dysregulation of ectodomain shedding is linked to diseases, including Alzheimer’s disease (AD) (Haass et al., 2012) and inflammation (McIlwain et al., 2012).

The physiological functions of ectodomain shedding are only beginning to be uncovered, particularly because the functions of most shed ectodomains are not known. For synaptic cell adhesion molecules (CAMs), proteolytic cleavage is generally thought to terminate adhesion, causing loss of cell-cell contacts and synapse weakening (Malinverno et al., 2010). Remarkably, ectodomains of some synaptic proteins participate in pathogenesis or have protective effects in disease conditions (Venkatesh et al., 2017; Yao et al., 2015). This suggests broader roles of synaptic ectodomains in pathogenic processes which remain to be explored.

The CSF is one of the few sources of proteins derived from the brain of living humans; hence it has the potential to offer a readout of brain physiopathology. Synaptic transmembrane proteins have been detected in the CSF (Pigoni et al., 2016), suggesting the CSF could be a source for biomarkers of synapse function or pathology in the brain. However, the relationship of the neuronal and, more specifically, the synaptic sheddome to the CSF has not yet been systematically investigated.

Disease-linked mutations in several synaptic membrane-anchored proteins predictably trigger a soluble secreted ectodomain. One such molecule is encoded by the *CNTNAP2* gene. Gene dosage, rare mutations, and common variations in *CNTNAP2* have been associated with several syndromal neurodevelopmental disorders sharing symptoms of epilepsy, intellectual disability (ID), ASD, and language impairment (Poot, 2015). The *CNTNAP2* gene encodes contactin associated protein like 2 (Caspr2 or CNTNAP2), which is abundant in the brain and it is enriched at synapses (Bakkaloglu et al., 2008). Moreover, CNTNAP2 also has a role in neuronal connectivity and synchrony. *CNTNAP2* polymorphisms disrupt brain connectivity (Scott-Van Zeeland et al., 2010). Synchronized network activity plays key roles during several stages of brain development, in sensory processing, motor control, and cognition (Uhlhaas and Singer, 2010). However, the molecular mechanisms regulating network synchrony are yet to be fully understood.

Here we hypothesized that proteomic analysis of the CSF and neuronal sheddomes could reveal novel synaptic regulators and functions. We showed that the neuronal sheddome is mirrored in hCSF and is enriched in adhesion and synapse regulatory molecules, as well as neurodevelopmental disorder risk molecules. We detected CNTNAP2 in hCSF and showed that it undergoes synaptic activity-dependent ectodomain shedding. We found that the ectodomain of CNTNAP2 (CNTNAP2-ecto) binds to and activates the ATP2B/PMCA Ca^2+^ pump and modulates neuronal network synchrony. Taken together, our data provide new insight into the biology of synaptic ectodomain shedding and reveal a novel mechanism of regulation of Ca^2+^ homeostasis and neuronal network synchrony.

## RESULTS

### The neuronal sheddome is mirrored in the CSF and is enriched in neurodevelopmental disorder risk factors

While proteomic analyses of cellular secretomes and sheddomes have been performed (Brown et al., 2013; Tien et al., 2017), the neuronal sheddome has not been systematically analyzed, and its relationship with the CSF proteome has not been investigated. While the term “secretome” refers to all proteins found in the extracellular milieu of a given biological sample, the “sheddome” is the subset of proteins found in the secretome that undergo ectodomain shedding and thus originally possess a transmembrane domain or a glycosylphosphatidylinositol (GPI) anchor.

To gain biological insight into the neuronal sheddome, we analyzed a previous dataset from secretome protein enrichment with click sugars and liquid chromatography tandem mass spectrometry (LC-MS/MS) analysis of media from cultured neurons. To determine the subset of proteins that undergo ectodomain shedding, we overlapped this dataset with proteins that are bound to the cell membrane (i.e., contain at least one transmembrane domain or a GPI anchor, based on UniProt). We identified 178 proteins likely to undergo ectodomain shedding, defined as the “neuronal sheddome” (Fig.1A). To gain insight into their biological functions, we performed Gene Ontology (GO) analysis. The most enriched biological processes were “positive regulation of synapse assembly” and “cell adhesion”.

**Figure 1.**
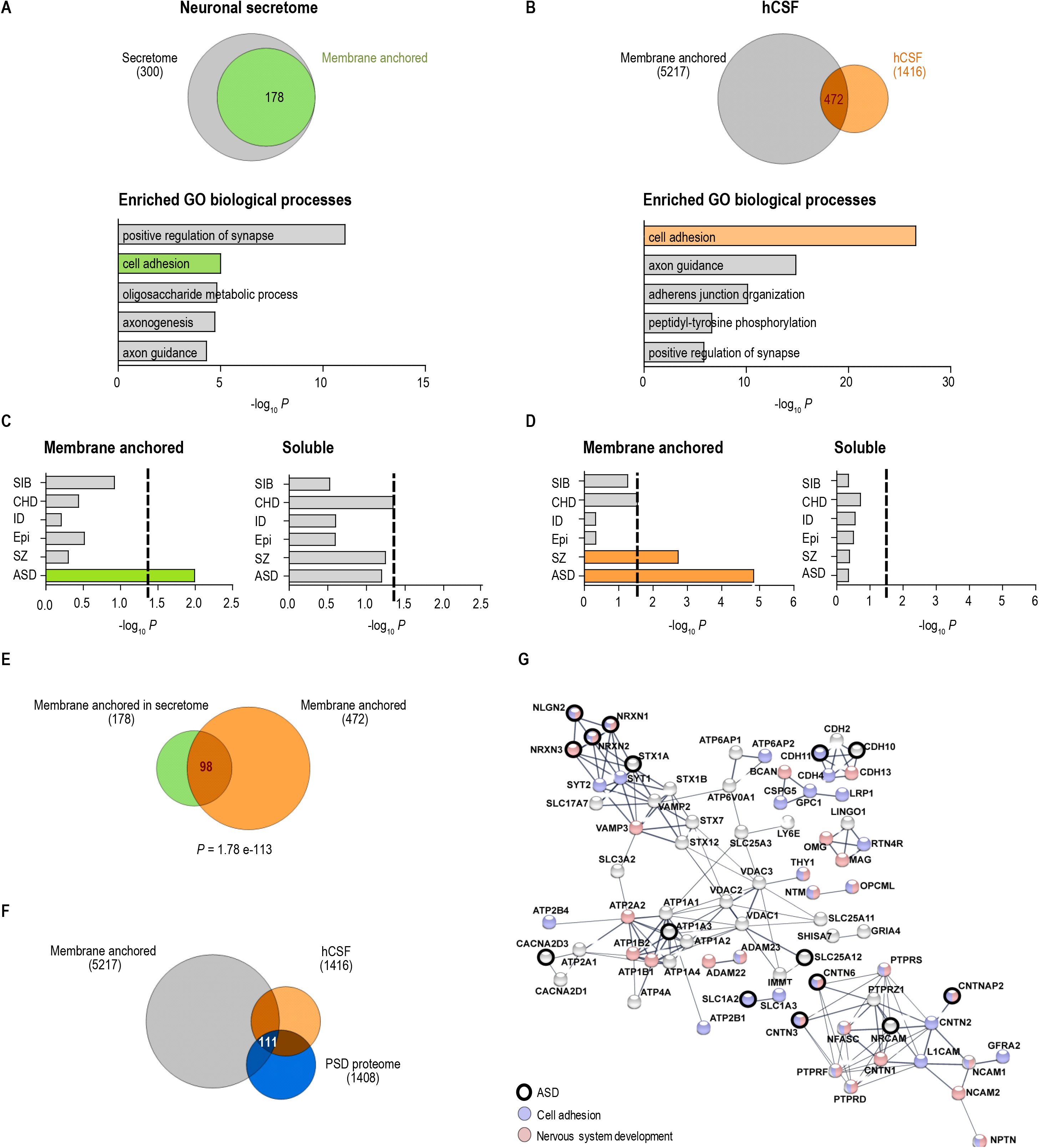
Bioinformatic analysis reveals the identity of the neuronal sheddome and shows an enrichment of ASD-related risk factors. **A.** Venn diagram of proteins found in the neuronal secretome (300) shows that 178 are membrane anchored (containing either a transmembrane domain or a GPI anchor) and are susceptible to ectodomain shedding. Gene ontology analysis of the *in vitro* neuronal sheddome shows an enrichment of proteins regulating synapse assembly or cell adhesion. **B.** Venn diagram showing that, of the 1416 proteins detected in human CSF, 472 potentially could undergo ectodomain shedding. GO analysis of membraned-anchored proteins shows cell adhesion and axon guidance as the top processes. **C.** Enrichment of disease-relevant genes in the neuronal secretome. Hypergeometric tests demonstrate an enrichment of proteins encoding disease-relevant genes in the membrane-anchored fraction but not the soluble fraction. **D.** The membrane-anchored fraction of the hCSF shows an enrichment for SZ- and ASD-related genes. **E.** Hypergeometric tests shows a highly significant overlap between the neuronal sheddome *in vitro* and the hCSF sheddome. **F.** Venn diagram depicting the “synaptic sheddome”. **G.** Protein-protein interaction network of the 111 proteins belonging to the synaptic sheddome. Proteins involved in cell adhesion processes are highlighted in blue, nervous system development is highlighted in red, and ASD-related genes according to SFARI are highlighted by bold circles. See also Figure S1.

To identify proteins in the CSF that potentially undergo ectodomain shedding, we performed LC-MS/MS analysis of human CSF (hCSF) and detected 1416 proteins. We overlapped it with a dataset of membrane-anchored proteins, using the same filtering method as for the neuronal secretome, yielding 472 proteins, the “CSF sheddome” (Fig.1B). GO analysis of these proteins revealed “cell adhesion” and “axon guidance” as the most significantly enriched biological processes. However, GO analysis of soluble proteins in the neuronal secretome and the hCSF showed an enrichment in “extracellular matrix organization” and “platelet degranulation” related proteins (Fig.S1D). When analyzed by number of proteins per biological process, cell adhesion was the most highly represented in the three datasets we analyzed: the neuronal sheddome, and two hCSF datasets (Fig.S1F).

Next, we examined the disease profiles of the *in vitro* neuronal secretome and the hCSF proteome. We overlapped either the membrane-anchored or the secreted proteins from both datasets with neurodevelopmental disorder-specific genes compiled from *de novo* exonic mutations, which implicate individual genes with particular disorders. Remarkably, proteins encoding ASD susceptibility genes were enriched in the neuronal sheddome, but not in the secreted components of the neuronal secretome (Fig.1C). Similarly, ASD and SZ risk gene-encoded proteins were enriched in the cleaved, but not in the secreted CSF fraction (Fig.1D). To corroborate the enrichment in ASD-related genes, we performed the same analysis using the gene list annotated by SFARI. We found again an enrichment for ASD-related molecules from SFARI in the membrane-anchored protein datasets, but not in the soluble protein datasets (Fig.S1E). This indicates that specifically ectodomain shedding, but not secretion of soluble proteins, may be implicated in ASD. We cross-validated these findings by performing GO and disease-profile analyses on an already published hCSF sheddome dataset with consistent results (Fig.S1A-B).

This analysis suggests that the CSF could mirror the neuronal sheddome, as they show similar GO and disease profiles. Corroborating this idea, hypergeometric testing showed a highly significant overlap between the neuronal and CSF sheddome datasets (Fig.1E and Fig.S1C). To specifically define the synaptic sheddome observable in the CSF, we overlapped the CSF sheddome with the postsynaptic density (PSD) proteome and found that 111 of 1408 PSD proteins (∼8%) were detected in the CSF sheddome (Fig.1F).

To gain insight into molecular and disease pathways within the synaptic sheddome present in hCSF, we built a protein-protein interaction (PPI) network (Fig.1G). This PPI network includes several neurodevelopmental disorder-associated synaptic molecules known to undergo ectodomain shedding (such as NRXN1/3, NLGN2), but also many proteins not yet reported to undergo ectodomain shedding, including CNTNAP2. This highlights the paucity of information on ectodomain shedding and suggests that analysis of the CSF could be a reasonable strategy to investigate synaptic ectodomain shedding in humans.

### CNTNAP2 undergoes activity-dependent ectodomain shedding

To further explore the hypothesis that the synaptic sheddome detectable in hCSF can reveal novel biological mechanisms, we focused our investigation on CNTNAP2, previously unknown to undergo ectodomain shedding.

We first validated the presence of CNTNAP2-ecto in hCSF and mouse CSF (mCSF) by western blotting (WB), using antibodies against the extracellular N-terminal (N-term) region and the intracellular C-terminal (C-term) region (Fig.2A). The N-term was present in both hCSF and mCSF, with a molecular weight just below that of the full-length protein containing the C-term region, suggesting cleavage of the protein to produce CNTNAP2-ecto (Fig.2B). The same band was also present in mouse cortex soluble fraction, but was absent from *Cntnap2* KO mouse brain homogenate (Fig.2C). The shorter N-term segment of CNTNAP2 was detected in whole cell lysates and extracellular media from WT but not *Cntnap2* KO mouse cortical neuronal cultures (Fig.2D). CNTNAP2 ectodomain shedding was also corroborated in HEK293 cells transfected with exogenous CNTNAP2 (Fig.S2A). Together, these data show that soluble CNTNAP2-ecto is present in CSF, in mouse brain, and *in vitro* in neuronal media.

**Figure 2.**
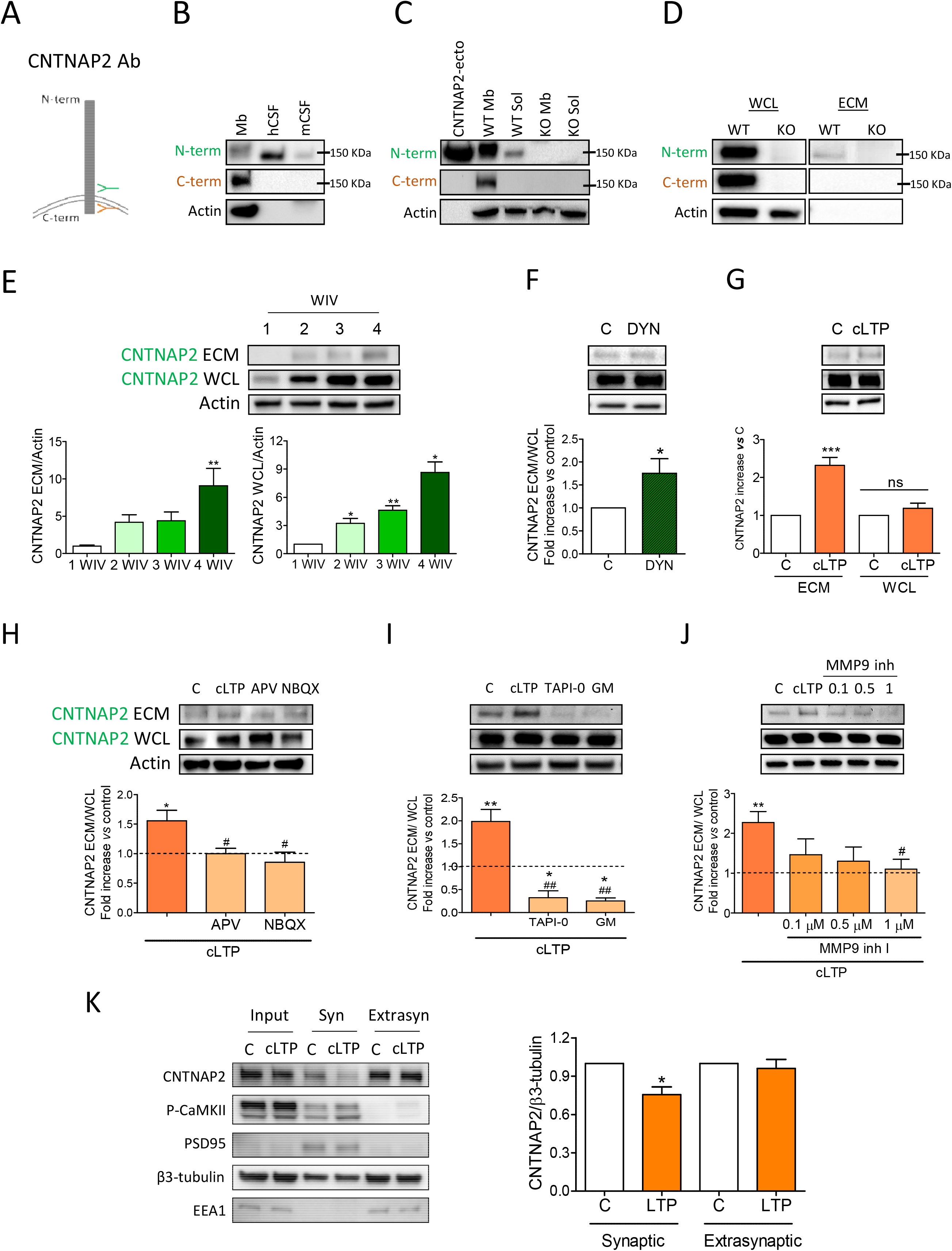
CNTNAP2 undergoes activity-dependent ectodomain shedding. **A.** Schematic showing the extracellular (green) and the intracellular (orange) antibodies against CNTNAP2. **B.** Western blot (WB) images of a membrane fraction (Mb), human CSF (hCSF), and mouse CSF (mCSF) using the N-term and the C-term antibodies against CNTNAP2. **C.** After fractionating *Cntnap2* WT and KO mouse brains to obtain membrane and soluble fractions, CNTNAP2-ecto is found in the soluble fractions of the WT but not of the KO. CNTNAP2-ecto produced in N2a cells is loaded as a positive control. **D.** WB images of *Cntnap2* WT and KO dissociated neurons show the presence of CNTNAP2 in the whole cell lysates (WCL) from WT neurons with both antibodies and signal in the extracellular media (ECM) with the N-terminal antibody. **E.** CNTNAP2 expression levels increase with time in culture in WCL (1, 2, 3, 4 weeks *in vitro* (WIV)) in dissociated rat neurons, in parallel to the levels of CNTNAP2-ecto levels in the ECM. Data are represented as mean±SEM. ECM data: n=3-7, Kruskal-Wallis test + Dunn’s *post hoc*, *P*=0.0061; WCL data: n=3-6, Mann-Whitney test, 1 *vs* 2 WIV *P*=0.0490, 1 *vs* 3 WIV *P*=0.012, 1 *vs* 4 WIV *P*=0.0168. **F.** Endocytosis inhibitor dynasore increases CNTNAP2 ectodomain shedding. Data are represented as mean±SEM. ECM data: n=13, Wilcoxon test, *P*=0.0398. **G.** cLTP increases the levels of CNTNAP2-ecto in the ECM with no changes being observed in WCL. Data are represented as mean±SEM. n=12, paired *t*-test, *P*<0.0001; WCL data: n=15, paired *t*-test, *P*=0.1780. **H.** APV and NBQX block cLTP-induced CNTNAP2 ectodomain shedding. Data are represented as mean±SEM. n=4-8; C *vs* cLTP paired *t*-test, \**P*<0.0167; cLTP *vs* NBQX/APV 2-way ANOVA *P*=0.0226 +Newman-Keuls *post hoc*, # *P*<0.05. **I.** Activity-dependent CNTNAP2 ectodomain shedding is prevented by the incubation of TAPI-0 and t GM6001. Data are represented as mean±SEM. n=7; C *vs* cLTP paired *t*-test, \*\**P*=0.0092; C *vs* TAPI Wilcoxon test, \**P*=0.0313, C *vs* GM Wilcoxon test, \**P*=0.015; cLTP *vs* TAPI/GM Friedman test *P*=0.0012 + Dunn’s *post hoc*, ## *P*<0.01. **J.** The specific MMP9 inhibitor I decreases CNTNAP2 ectodomain shedding upon cLTP induction in a concentration-dependent manner. Data are represented as mean±SEM. n=5-9; C *vs* cLTP paired *t*-test, \*\**P*<0.0016; cLTP *vs* MMP9 inhibitor I Kruskal-Wallis *P*=0.022 + Dunn’s *post hoc*, # *P*<0.05. **K.** Subcellular fractionation of rat neurons shows that the levels of N-term CNTNAP2 decrease in the synaptic fraction after cLTP. Data are represented as mean±SEM. n=3-4; Synaptic C *vs* cLTP paired *t*-test, \**P*=0.026; extrasynaptic C *vs* cLTP paired *t*-test, *P*=0.62.

CNTNAP2-ecto levels increased in media of cultured neurons from week 1 to 4, as neurons and synapses mature (Fig.2E). The amount of CNTNAP2-ecto in the media paralleled that of full-length CNTNAP2 in whole cell lysates, indicating that its increased shedding was due to increased overall protein levels.

We then asked whether CNTNAP2 was cleaved at the plasma membrane. Incubation of rat neurons with the endocytosis inhibitor dynasore significantly increased CNTNAP2-ecto in the media. As inhibiting endocytosis increases surface levels of plasma membrane proteins, this indicates that CNTNAP2 cleavage occurs at the cell surface (Fig.2F).

Because cleavage of several other synaptic adhesion molecules is regulated by synaptic activity (Suzuki et al., 2012), we examined the effect of chemical LTP (cLTP) on CNTNAP2-ecto production. cLTP of week 4 neuronal cultures resulted in a strong increase in CNTNAP2-ecto in the neuronal media (Fig.2G), indicating that CNTNAP2-ecto shedding is up-regulated during synaptic potentiation. cLTP-dependent ectodomain shedding required both NMDA and AMPA receptor activity, as it was blocked by both aminophosphonovalerate (APV) and NBQX (Fig.2H).

We next used different protease inhibitors to identify the protease responsible for CNTNAP2 cleavage. cLTP-induced CNTNAP2 ectodomain shedding was blocked by TAPI-0, which inhibits several MMPs and TACE (Fig.2I). In addition, the general MMP inhibitor GM6001 significantly reduced cLTP-induced CNTNAP2 cleavage (Fig.2I), implicating MMPs in activity-dependent CNTNAP2 cleavage. As only MMP3 and 9 are highly expressed in cortex, and MMP9’s activity is known to be up-regulated by LTP, we performed a dose-response curve using MMP9 inhibitor I. This inhibitor blocked cLTP-dependent CNTNAP2-ecto shedding in a concentration-dependent manner (Fig.2J), indicating that MMP9 mediates activity-dependent CNTNAP2-ecto shedding. These results were validated in CNTNAP2-transfected N2a cells, where GM6001, TAPI-0 and MMP9 inhibitor inhibited CNTNAP2 ectodomain shedding (Fig.S2B).

We then tested whether activity-induced CNTNAP2 cleavage occurs at synaptic sites by subcellular fractionation. We observed a decrease of CNTNAP2 levels in the synaptic fraction after cLTP and no change in the extrasynaptic fraction (Fig.2K), indicating that CNTNAP2 occurs in the synaptic region.

### Spatial organization of CNTNAP2 ectodomain shedding

To examine the distribution of CNTNAP2-ecto *in situ*, we sparsely transfected cortical neurons generated from *Cntnap2* KO mice with a distal N-terminally flag-tagged CNTNAP2 along with GFP (Fig.3A). We detected abundant exogenous Flag-CNTNAP2 in soma of individual neurons, as well as in dendrites and spines (Fig.3A-B). Flag-CNTNAP2 immunofluorescence could also be detected outside the neurons, not overlapping with GFP, as shown by line scans (Fig.3B).

**Figure 3.**
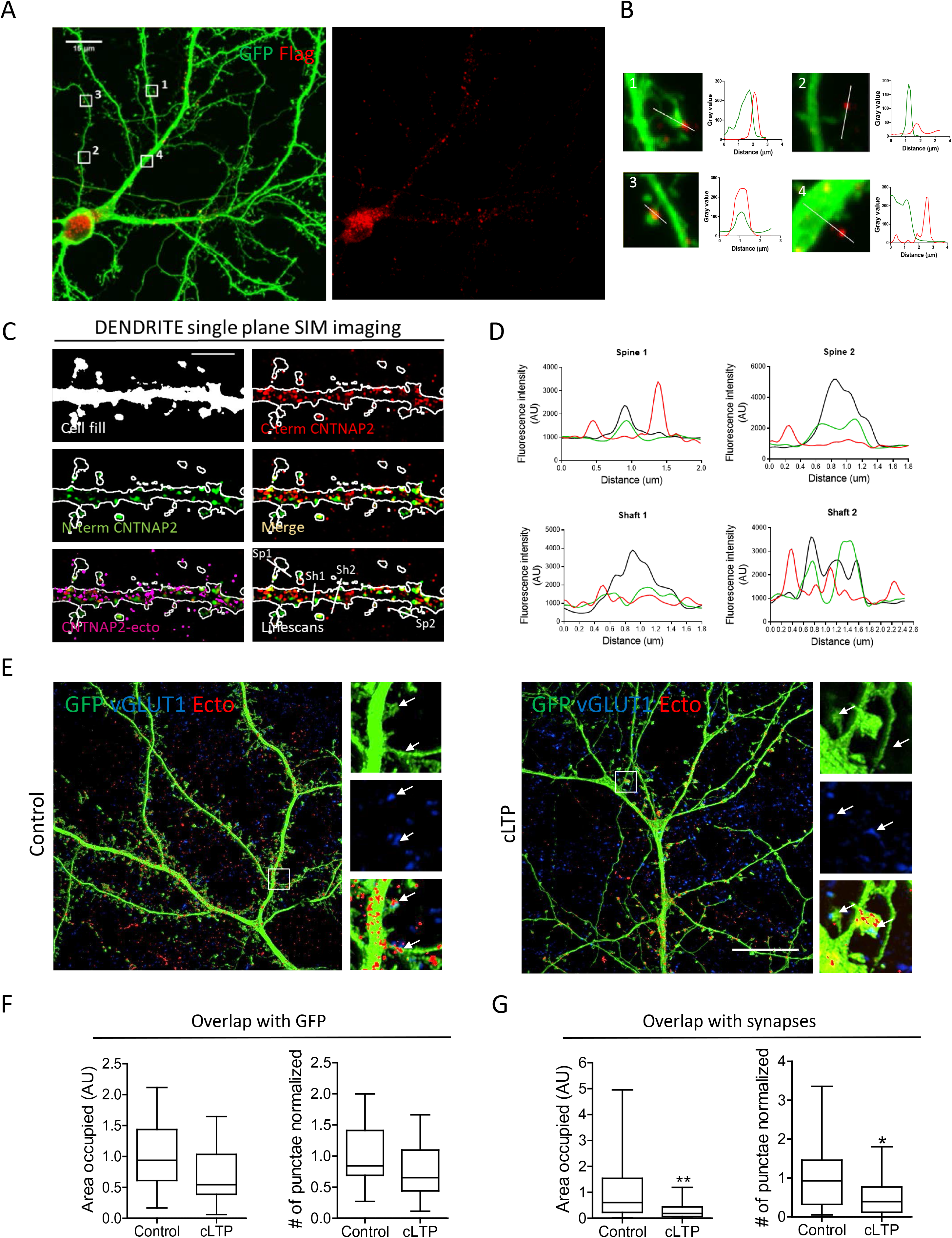
Spatial organization of CNTNAP2 ectodomain shedding. **A.** Confocal microscopy images of *Cntnap2* KO neurons transfected with Flag-CNTNAP2 and GFP and stained for both. Scale bar 15 μm. **B.** Line scans show the presence of the CNTNAP2 extracellularly close to a dendritic spine (1), outside the shaft (2, 4), and surrounding a spine neck (3). **C-D.** SIM imaging and line scan analysis of CNTNAP2-ecto in the dendritic compartment. Scale bar 5 μm. **E.** SIM image analysis without (left) or with (right) cLTP. Scale bar 15 μm. **F-G.** The area occupied by CNTNAP2-ecto and the number of punctae in the GFP compartment (F) and in the synaptic regions (G). Data are presented as median and quartiles, n=20-22 neurons, N=3 cultures, 2 coverslips/condition/culture. Area occupied by CNTNAP2-ecto in GFP, Mann-Whitney test, *P*=0.12; # of punctae in GFP, *t*-test, *P*=0.1632. Area occupied by CNTNAP2-ecto in synapses, Mann-Whitney test, \*\**P*=0.006; # of punctae in synapses, Mann-Whitney test, \**P*=0.024. See also Figure S3.

To further enhance the spatial resolution, we used structured illumination microscopy (SIM) to visualize the relative distribution of the N- and C-termini of transfected Flag-CNTNAP2 in GFP-expressing neurons generated from *Cntnap2* KO mice (Fig.3C). In total, 55% of CNTNAP2-ecto was found within the GFP signal and the rest was found outside (Fig.S3A). CNTNAP2 N- and C-terminal immunofluorescence colocalized inside dendrites and spines, but not outside, and N-terminal immunofluorescence was detected outside neurons, often near spines and dendrites (Fig.3C) as well as axons and synaptic boutons (Fig.S3B), consistent with it being cleaved and deposited in the extracellular space surrounding neurons.

To examine the spatial organization of activity-dependent CNTNAP2 ectodomain shedding, we used SIM to visualize the distribution of the N- and the C-termini of CNTNAP2 in GFP-expressing *Cntnap2* KO neurons subjected to cLTP (Fig.3D). We analyzed the overlap of CNTNAP2-ecto (presence of N-termini but absence of C-termini) relative to the neuronal structure, outlined by GFP fluorescence. We found no difference in the area occupied or the number of CNTNAP2-ecto puncta within the GFP signal between control and cLTP-treated neurons (Fig.3E-F). However, analyzing the same parameters in the synaptic compartment, defined as the overlap between GFP and the presynaptic marker vGLUT1, we found that upon cLTP, both the area occupied and the number of CNTNAP2-ecto of puncta significantly decreased in the synaptic region (Fig.3E, G). These data suggest that CNTNAP2-ecto, produced postsynaptically upon cLTP, diffuses away from synaptic regions, likely in a paracrine fashion.

### CNTNAP2-ecto affinity-MS identifies PMCA2 as a novel interacting partner

Notably, only two ligands of CNTNAP2-extracellular domain are known, CNTN1 and CNTN2 (Poliak et al., 2003; Rubio-Marrero et al., 2016). Because a few reports also identified signaling roles for ectodomains, we hypothesized that shed CNTNAP2-ecto may act as a ligand for unidentified transmembrane proteins. We used a combination of CNTNAP2-ecto affinity chromatography with a proteomic protocol that couples affinity purification with LC-MS/MS and bioinformatics analysis (Fig.4A). We generated a recombinant flag-tagged CNTNAP2-ecto bait protein from the supernatants of CNTNAP2-transfected HEK293 cell cultures. SYPRO ruby staining of the recombinant CNTNAP2-ecto and vehicle control revealed high purity of the purified CNTNAP2-ecto (Fig.4B). Moreover, MS analysis of tryptic peptide digestion of the purified CNTNAP2-ecto identified peptides corresponding only to the extracellular domain, indicating little or no contamination with full-length protein (Fig.4C). To identify proteins interacting with the CNTNAP2-ecto, we incubated the Flag-CNTNAP2-ecto or vehicle control with mouse cortical membranes. Affinity-purified proteins were comprehensively analyzed by LC-MS/MS. To sort the identified proteins, we rank-ordered the prey proteins detected in descending order on the basis of the number of normalized spectral counts. The fold enrichment of each prey protein was calculated by subtracting the number of spectral counts in the sample containing the bait minus the sample containing the vehicle control, this value was then divided by the total number of spectral counts obtained for CNTNAP2-ecto. As expected, the bait protein was the most abundant protein present. We also detected CNTN1 as a very abundant affinity-isolated protein, validating our approach (Fig.4D). In addition, all four members of the PMCA/ATP2B family of Ca^2+^ transporters were highly abundant among CNTNAP2-ecto interacting proteins (Fig.4E). Because PMCA2 showed a high enrichment and because it has been implicated in ASD (Takata et al., 2018), we investigated its interaction with CNTNAP2-ecto.

**Figure 4.**
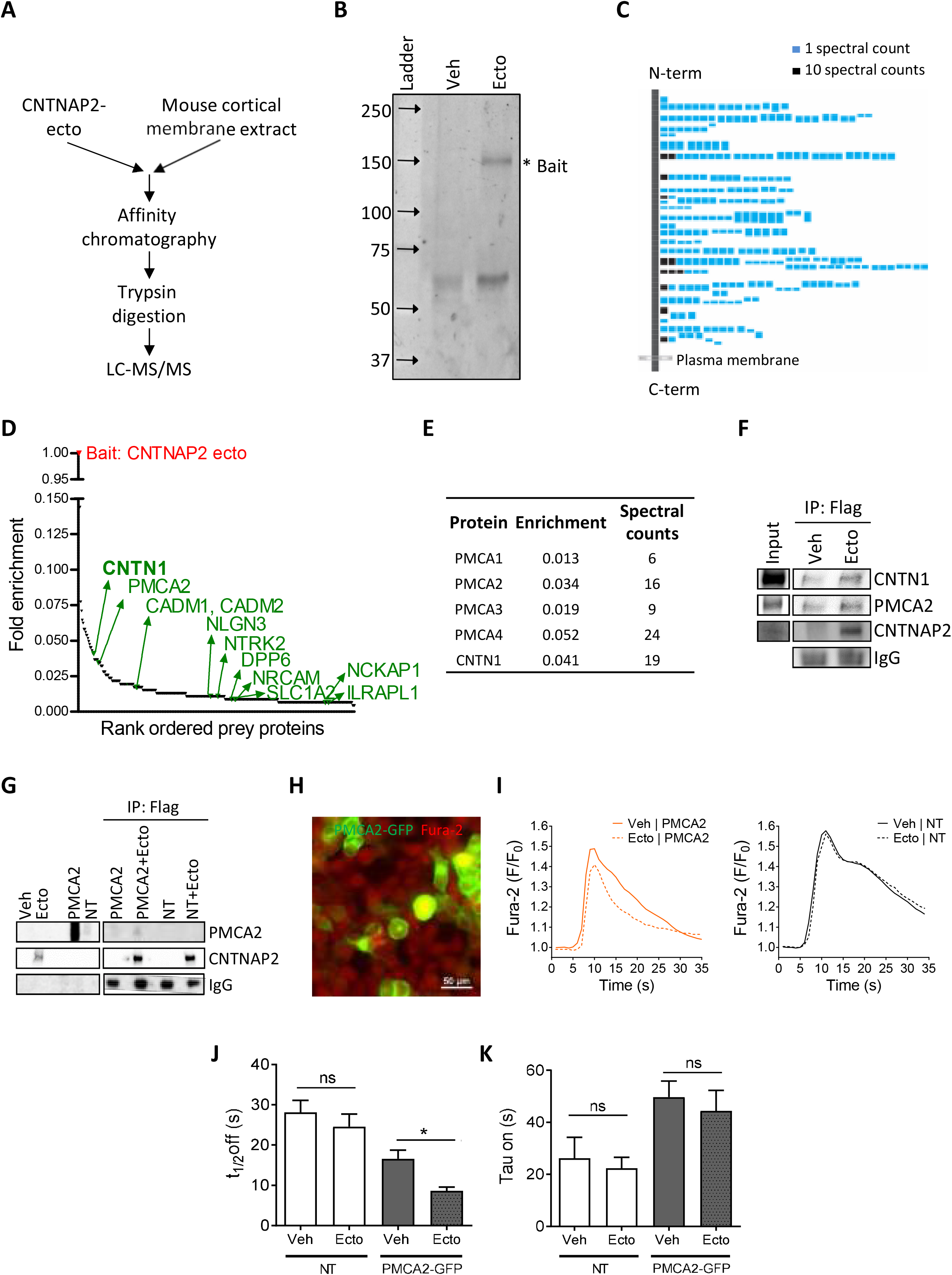
Pulldown coupled to LC-MS/MS identifies CNTNAP2-ecto as a binding partner and activator of PMCA2. **A.** Schematic of the process followed. Membranes from mice cortices were incubated with CNTNAP2-ecto and subjected to affinity chromatography. After trypsin digestion of the pull-down, samples were analyzed by LC-MS/MS. **B.** SYPRO Ruby staining of the CNTNAP2-ecto band right below 150 kDa. Only one non-specific band is found between 50-75 kDa, probably corresponding to BSA. **C.** Mapping of the peptides found in the CNTNAP2 fragment. **D.** Fold enrichment of rank-ordered prey proteins detected by LC-MS/MS highlighting ASD-related risk factors and the positive control contactin-1 in green. **E.** Fold enrichment and spectral counts of the 4 isoforms of PMCA and CNTN1 detected by LC-MS/MS. **F.** Confirmation of interaction between CNTNAP2-ecto and PMCA2 in mouse cortical membranes by WB. **G.** WB confirming the interaction between CNTNAP2-ecto and PMCA2 in HEK293 cells transfected with PMCA2. Cell lysates were incubated with CNTNAP2-ecto, and flag agarose beads were used to pull down CNTNAP2-ecto. **H.** Epifluorescence microphotography of HEK cells transfected with PMCA2-GFP and loaded with Fura2 (red). **I.** Ca^2+^ curve elicited by ATP in PMCA2-transfected or non-transfected (NT) HEK293 cells in the presence or absence of CNTNAP2-ecto. **J.** t_1/2_ off analysis of the Ca^2+^ curve elicited by ATP in the presence or absence of CNTNAP2-ecto in transfected and NT HEK cells showed a specific effect of CNTNAP2-ecto only on PMCA2-positive cells. Data are represented as mean±SEM, N=6 cultures, n=13 dishes, one-way ANOVA, \*\*\**P*<0.0001; Newman-Keuls *post-hoc*, \**P*<0.05. **K.** Comparison of the tau_on_ data shows no differences between groups on the ascending phase of the Ca^2+^ curve elicited by ATP. Data are represented as mean±SEM, N=6 cultures, n=13 dishes, Kruskal-Wallis test, \*\*\**P*=0.0004; Dunn’s *post-hoc* ^ns^*P*>0.05. See also Figure S4.

To validate the interaction, we applied purified CNTNAP2-ecto or vehicle control to mouse cortical membrane fractions, and used flag-agarose beads to pull down CNTNAP2-ecto, followed by detection of co-purified PMCA2 by WB (Fig.4F). To corroborate this finding, we incubated purified WT and 1253* CNTNAP2-ecto or vehicle with lysates of PMCA2-transfected or untransfected HEK293 cells, and detected the interaction by WB (Fig.4G and Fig.S4A). Moreover, confocal images of dissociated rat cortical neurons showed colocalization of PMCA2 and CNTNAP2 (Fig.S4B). To determine if absence of CNTNAP2 impacted the developmental time course of PMCA2 expression, we compared the protein levels of PMCA2 in WT and *Cntnap2* KO mouse cortex and found a significant reduction in PMCA2 levels in the KO cortex at P28 (Fig.S4C). These experiments suggest that PMCA2 is a novel endogenous CNTNAP2-ecto binding partner.

### CNTNAP2-ecto activates Ca^2+^ extrusion in neurons and acute brain slices

PMCA2 regulates cytosolic Ca^2+^ by extruding it into the extracellular space with the concomitant hydrolysis of ATP. To investigate directly whether CNTNAP2-ecto regulates the Ca^2+^ pump activity of PMCA2, we expressed exogenous PMCA2 in HEK293 cells, loaded them with the fluorescent Ca^2+^ sensitive dye Fura-2, and incubated them with CNTNAP2-ecto or vehicle control for 5 min (Fig.4H). Immediately after, we treated the cultures with 0.5 mM ATP to elicit a cytosolic Ca^2+^ signal (Fig.4I and Fig.S4G). CNTNAP2-ecto reduced the t_1/2_ off of the Ca^2+^ peak by ∼ 50%, without affecting non-transfected HEK293 cells (Fig.4J, K). However, neither the Ca^2+^ peak amplitude (Fig.S4G) nor the tau_on_ (Fig.4J) were affected by the CNTNAP2-ecto. This indicates that CNTNAP2-ecto specifically increases the speed of Ca^2+^ extrusion by PMCA2.

We then assessed the effect of CNTNAP2-ecto on neurons transfected with the Ca^2+^ indicator GCamP6s and the cell fill DsRedExpress2 to check for cell health (Fig.5A). We first performed a concentration response curve, finding that 10 nM is the CNTNAP2-ecto concentration exerting the strongest effect in Ca^2+^ extrusion (Fig.S5B). Incubation with CNTNAP2-ecto for 5 min decreased by 30% the t_1/2_ off of the KCl-induced Ca^2+^ signal (Fig.5A-C), without affecting the tau_on_ or the amplitude (Fig.5C and Fig.S5C). We then checked if the observed effect is specific to inhibitory or pyramidal neurons, finding that the most consistent effect occurs in excitatory neurons (Fig.S5D). These data indicate that CNTNAP2-ecto specifically regulates the extrusion phase of the Ca^2+^ signal in primary neurons.

**Figure 5.**
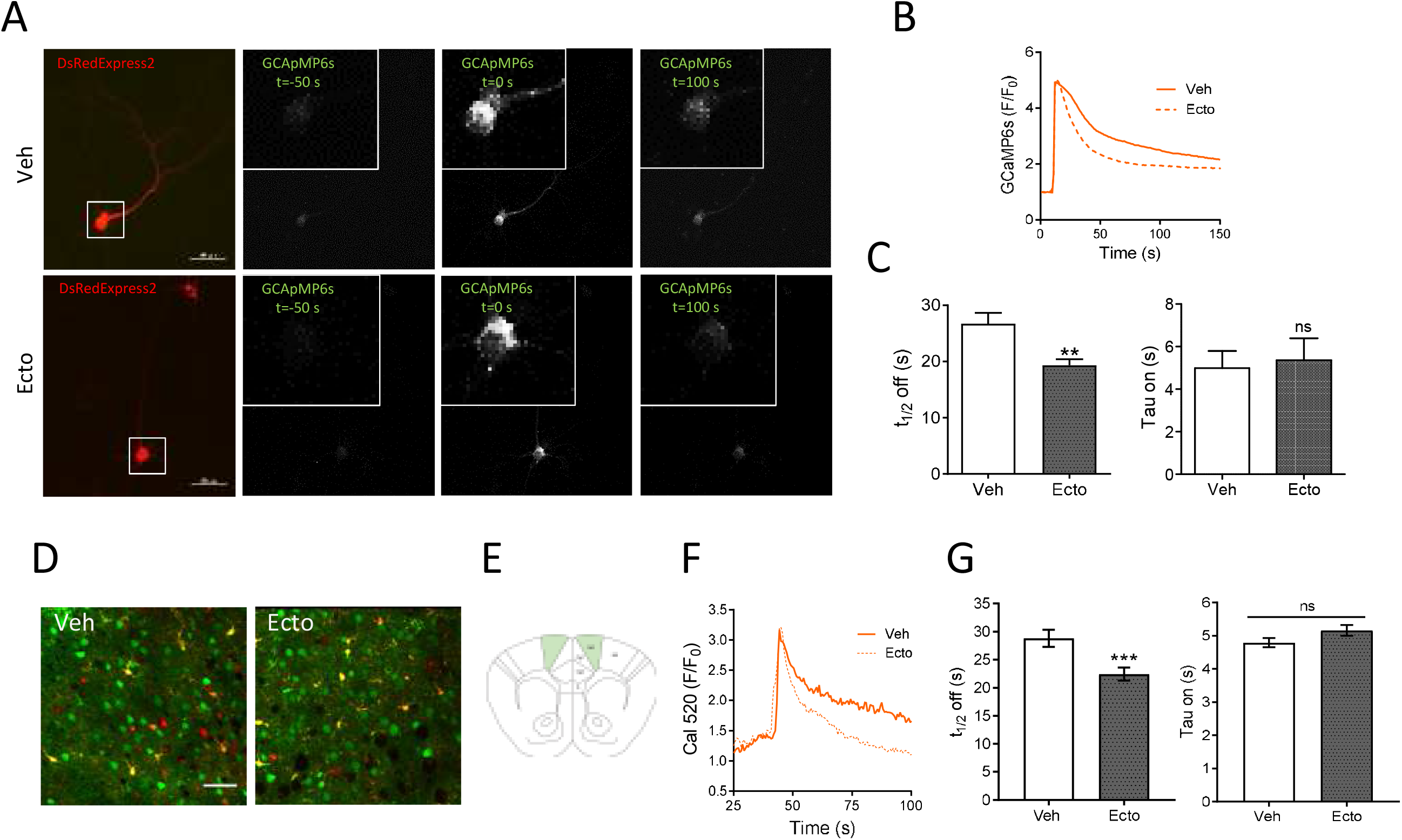
CNTNAP2-ecto activates Ca^2+^ extrusion in dissociated rat cortical neurons and in brain slices. **A.** Epiflourescence microphotographs of dissociated rat neurons transfected with the Ca^2+^ indicator GCaMP6s and the cell fill DsRedExpress2 at different time points before and after incubation with 30 mM KCl. **B.** Curve depicting the Ca^2+^ peak elicited by KCl in rat neurons after treatment with CNTNAP2-ecto or vehicle. **C.** Bar graph of the Ca^2+^ t_1/2_ off and tau_on_. Data are represented as mean±SEM, N=5 cultures, n=35/28 neurons, t_1/2_ off Mann-Whitney test, \*\**P*=0.0027; tau_on_ Mann-Whitney test, ^ns^ *P*=0.7651. **D.** Multiphoton microphotographs showing Cal520 staining in green, for recording cytosolic Ca^2+^ changes, and the glial indicator SR-101 in red. Scale bar=50 μm. **E.** Area imaged in acute brain slices obtained from CD1 mice. **F.** Ca^2+^ curve in brain slices treated with vehicle or CNTNAP2-ecto. **G.** Analysis of the Ca^2+^ curves shows a decrease in the t_1/2_ off but no effect on the tau_on_. Data are represented as mean±SEM, t_1/2_ off n=140/153 neurons, Mann-Whitney test, \*\*\**P*=0.0004; tau_on_ n=117/116 neurons, Mann-Whitney test, ^ns^*P*=0.058. See also Figure S5.

To investigate the effect of CNTNAP2-ecto in the context of native tissue with preserved cellular populations and networks, we measured Ca^2+^ dynamics after KCl-evoked Ca^2+^ responses in acute brain slices, imaging in layer V of the M2 region of the cortex (Fig.5D and E). We loaded slices with the cytosolic Ca^2+^ sensor Cal520-AM and the glial red dye SR-101 to discard the signal originating from glial cells, incubated them with CNTNAP2-ecto (Fig.5D), and imaged KCl-evoked Ca^2+^ signals (SuppVideo1). As in cultured neurons and HEK293 cells, CNTNAP2-ecto decreased the t_1/2_ off by 22% without affecting the tau_on_ and the Ca^2+^ amplitude (Fig.5F-G). CNTNAP2-ecto thus enhances Ca^2+^ extrusion in acute brain slices.

### CNTNAP2-ecto regulates synchrony in neuronal networks *in vitro* and in acute brain slices

In order to understand the impact that this newly discovered effect of CNTNAP2-ecto on Ca^2+^ homeostasis could have at the circuit level, we investigated how exposure to exogenous CNTNAP2-ecto affected coordinated patterns of Ca^2+^ signaling or Ca^2+^ synchrony. We first examined the effect of CNTNAP2-ecto on network activity in rat neurons *in vitro* by imaging the spontaneous Ca^2+^ transients or oscillations of cytoplasmic [Ca^2+^], using the Ca^2+^ dye Fluo-4 (Fig.6A). After normalizing the average somatic Ca^2+^ fluorescence for hundreds of neurons per coverslip, Ca^2+^ transients were automatically detected/defined in an unbiased manner using a custom-made Matlab routine (Fig.S6). Both vehicle- and CNTNAP2-ecto-treated neurons displayed synchronized network events (Fig.6B); however, CNTNAP2-ecto incubation led to a 25% decrease in the fraction of synchronized transients (Fig.6C, D), without changing the transient frequency (Fig.6E). By measuring the amplitude of the Ca^2+^ peaks within and outside of the synchrony windows, we observed an increase in the Ca^2+^ peak amplitude during network synchrony that was lost after CNTNAP2-ecto exposure (Fig.6F). This indicates that CNTNAP2-ecto suppresses the synchrony of Ca^2+^ firing and as a result, reduces the potentiation on the Ca^2+^ peak amplitude that occurs in synchronized events at a network level.

**Figure 6.**
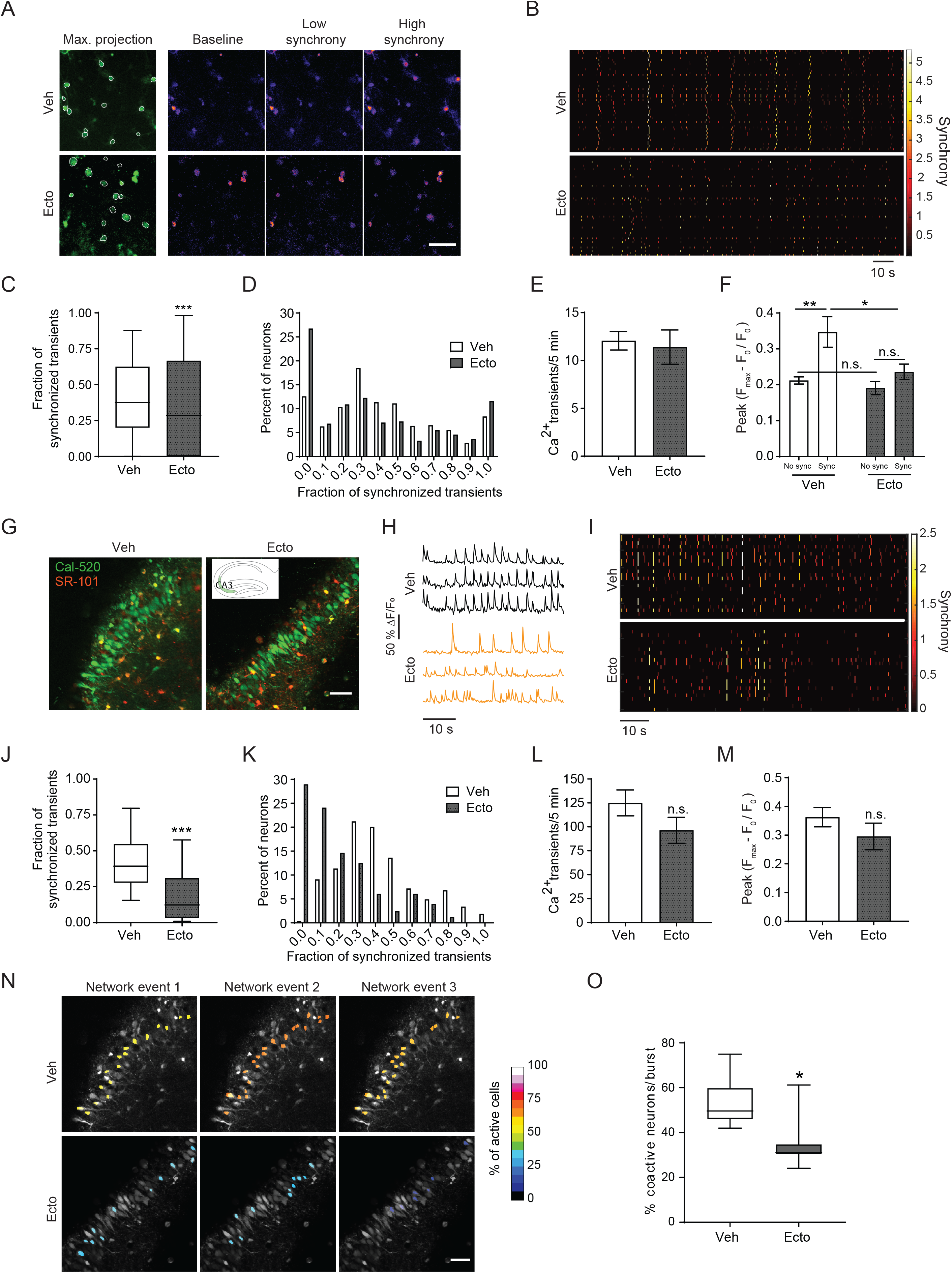
CNTNAP2-ecto regulates neuronal network synchrony *in vitro* and *ex vivo.* **A.** Microphotographs showing Fluo4 staining in dissociated rat neurons and heat maps depicting Ca^2+^ levels at baseline, low synchrony, and high synchrony states. Scale bar=50 μm. **B.** Raster plot depicting Ca^2+^ peaks color coded for the degree of network synchrony (1 equals the threshold of significance) shows a decrease in synchrony after treatment with CNTNAP2-ecto. **C.** Analysis of synchronized firing shows a decrease in CNTNAP2-ecto treated cells. Data are represented as median and quartiles, n=810/871 neurons, Mann-Whitney test, \*\*\**P*<0.0001. **D.** Histogram representing the distribution of synchronized firings per neuron shows an increase of the percentage of asynchronous neurons after CNTNAP2-ecto. **E.** Frequency of Ca^2+^ transients did not change. Data are represented as mean±SEM, n=7/8, *t*-test, *P*=0.75. **F.** Analysis of the peak of the Ca^2+^ bursts shows a decrease only in the synchronous Ca^2+^ peaks after CNTNAP2-ecto treatment but not in the asynchronous peaks. Data are represented as mean±SEM, n=7-8 slices, 2-way ANOVA test, interaction *P*=0.088, row factor (synchrony) *P*=0.0015, column factor (treatment) *P*=0.0149. Bonferroni *post-hoc* test \*\**P*=0.007, \**P*=0.028. **G.** Brain slices stained with the Ca^2+^ indicator Cal520 (green) and the glial dye SR-101 (red) at the CA3 region of the hippocampus. Scale bar=50 μm. **H.** Ca^2+^ spontaneous signals taken from three different neurons from vehicle-treated (black) and CNTNAP2-ecto-treated (orange) slices. **I.** Raster plot depicting Ca^2+^ bursts in time in hippocampal slices shows a decrease in synchrony after treatment with CNTNAP2-ecto. **J.** Quantification of Ca^2+^ dynamics shows a decrease in the synchronized firing in CNTNAP2-ecto treated cells. Data are represented as median and quartiles, n=264/328 neurons, Mann-Whitney test, \*\*\**P*<0.0001. **K.** Histogram representing the frequency of neuronal Ca^2+^ synchrony shows an increase in the fraction of asynchronous and low synchronous neurons(synchrony ratio 0-0.2) in CNTNAP2-ecto treated cells. **L.** Analysis of Ca^2+^ firing frequency in vehicle and CNTNAP2-ecto treated slices. Data are represented as mean±SEM, n=7/8, *t*-test, *P*=0.16. **M.** Quantification of the amplitude of the Ca^2+^ peaks. Data are represented as mean±SEM, n=7/8, *t*-test, *P*=0.25. **N.** Representative images of three network events occurring in the same slice in either vehicle-treated or CNTNAP2-ecto-treated conditions, showing the % of active cells in an individual Ca^2+^ burst of activity. Scale bar 50 μm. **O.** Quantification of the percentage of active neurons during a Ca^2+^ burst of activity after vehicle or CNTNAP2-ecto treatment. Data are represented as median and quartiles, n=810/871 neurons, Mann-Whitney test, \**P*=0.014.

To investigate whether CNTNAP2-ecto could also alter neuronal Ca^2+^ patterns in a preparation with intact circuitry, we examined spontaneous neuronal activity in acute brain slices from P5-P7 mice (Fig.6G-O). We imaged network events by two-photon microscopy in the CA3 region of the hippocampus with bath application of modified artificial cerebrospinal fluid (aCSF). In agreement with previous studies (Bonifazi et al., 2009), vehicle-treated slices exhibited regular network events (Fig.6H-I, SuppVideo2). Remarkably, CNTNAP2-ecto incubation decreased the fraction of peaks localized inside these synchronized events by 30% (Fig.6J-K). This difference was not associated with any significant changes in Ca^2+^ signal frequency or amplitude (Fig.6L-M). A fundamental aspect of neuronal networks is their ability to form functional assemblies of co-active cells, essential for cognitive processes and response to external stimuli. We analyzed these ensembles by measuring the percentage of co-active neurons during synchronized network events and, interestingly, these were reduced by 34.3% when treated with CNTNAP2-ecto (Fig.6N, O). In summary, CNTNAP2-ecto alters the neuronal network behavior in brain slices by desynchronizing neuronal firing.

## DISCUSSION

### The neuronal sheddome is mirrored in the CSF and is enriched in neurodevelopmental disorder risk factors

Here we present a global bioinformatics characterization of the neuronal sheddome and show that it is closely mirrored in the CSF. We found that both the neuronal sheddome and CSF are enriched in adhesion and synaptic proteins, as well as in neurodevelopmental disorder risk molecules. While ectodomain shedding by individual synaptic molecules has been reported, the sheddome comprised of synaptic proteins has not been systematically analyzed, and its relationship with the CSF proteome, as well as the representation of neurodevelopmental disorder risk factors in the synaptic sheddome, have not been investigated. Our findings thus provide novel insight into the neurobiology of ectodomain shedding.

### Neurodevelopmental disorder risk factor CNTNAP2 undergoes synaptic activity-dependent ectodomain shedding

Proteomic profiling of the rodent neuronal sheddome *in vitro* and hCSF revealed that the prominent neurodevelopmental disorder risk factor CNTNAP2 undergoes synaptic activity-dependent ectodomain shedding mediated by MMP9, and it can be detected in hCSF. CNTNAP2 ectodomain shedding has not been reported before and may represent a mechanism of intercellular signaling. This indicates that proteolytically-cleaved cell-cell adhesion membrane proteins could function as signaling molecules.

SIM microscopy showed that CNTNAP2-ecto was often present outside of the neurons, sometimes at a considerable distance. Upon cLTP we observed a decrease in the amount of CNTNAP2-ecto in synaptic regions, similar to what was observed biochemically, consistent with ectodomain shedding occurring at synaptic sites. Notably, a recent article reported that postynaptically-released neurexin-1 binds to presynaptic UNC-2/CaV2 Ca^2+^ channels and reduces acetylcholine release at the neuromuscular juction (Tong et al., 2017).

### PMCA2 is a novel binding partner of CNTNAP2-ecto

Using affinity purification combined with MS proteomics, we identified novel targets of CNTNAP2-ecto, including the Ca^2+^ pump PMCA2. Currently, only two extracellular CNTNAP2 ligands are known, CNTN-1 and -2 (Poliak et al., 2003; Rubio-Marrero et al., 2016). Interaction of CNTNAP2-ecto with several PMCAs indicates that Ca^2+^ signaling is a major target of CNTNAP2 ectodomain signaling, and reveals a novel mechanism for the regulation of Ca^2+^ homeostasis by shed ectodomains.

Accurate control of intracellular Ca^2+^ is crucial for neuronal function. The plasma membrane Ca^2+^-ATPase (PMCA) family of Ca^2+^ pumps are the major high-affinity Ca^2+^ extrusion pathway in spines (Scheuss et al., 2006). CNTNAP2-ecto enhances Ca^2+^ extrusion through PMCAs, and may thus control local or global Ca^2+^ transients by shaping their duration and spread. Among PMCAs, PMCA2 is widely distributed throughout the brain (Fig.S4D) and is present both pre- and postsynaptically (Burette and Weinberg, 2007).

### A shed ectodomain regulates neuronal Ca^2+^ spikes and network synchrony

We show for the first time that neuronal synchrony is regulated by a shed ectodomain, through a novel mechanism of regulation of Ca^2+^ dynamics. Neural network synchrony plays key roles during several stages of brain development (Marin, 2016), in sensory processing, motor control, and cognition (Uhlhaas and Singer, 2010).

A brief activity-regulated increase of CNTNAP2-ecto could mediate feedback inhibition in response to a short period of high activity, such as following LTP, to reestablish normal activity levels, to prevent over-excitation and excessive synchrony. Alternatively, it may modulate activity-dependent shaping of oscillations, for example during sensory processing. Consistent with this, *Cntnap2* KO mice show altered network activity and have seizures (Penagarikano et al., 2011).

Mechanistically, we propose that CNTNAP2-ecto activates PMCA pumps and enhances cellular Ca^2+^ extrusion. Remarkably, even moderate changes in Ca^2+^ kinetics are known to have an impact on the entire neuronal network (Lavi et al., 2014).

### The synaptic sheddome as a potential source of biomarkers and players in pathogenesis

We show that the CSF mirrors the neuronal sheddome, potentially providing access to the extrasynaptic milieu in the brain. CSF could provide insight into normal and pathophysiological processes in the brain, and has been proposed as a source of novel biomarkers. For example, in neurodegenerative disorders including ALS, Parkinson’s, and AD, elevated levels of synaptic proteins in the CSF is thought to reflect synapse loss (Bereczki et al., 2017; Kester et al., 2015). Similarly, elevated levels of L1, NCAM, and p75NTR ectodomains have been detected in the CSF in AD (Jiao et al., 2015; Strekalova et al., 2006), ICAM5 in acute encephalitis (Lindsberg et al., 2002), and NCAM in SZ (Vawter et al., 2001). In contrast, because its shedding is activity-dependent, altered levels of CNTNAP2-ecto in the CSF could reflect alterations in activity in brain regions enriched in CNTNAP2.

Ectodomain shedding can be affected by disease-associated mutations, providing a new pathogenic mechanism for neurodevelopmental disorders. Of particular interest is the presence of the same homozygous truncating point mutation (I1253X) in several members of an old-order Amish population with cortical dysplasia focal epilepsy (CDFE), along with learning, language, and social deficits. In a cellular system, this mutation results in a secreted protein with a sequence nearly identical to native CNTNAP2-ecto (Falivelli et al., 2012).

Some of the synaptic protein ectodomains we have detected may participate in pathogenesis or have therapeutic utility. For example, activity-dependent shedding of neuroligin-3 promotes glioma proliferation (Venkatesh et al., 2017). Similarly, excessive CNTNAP2-ecto could promote pathogenesis by desynchronizing networks. On the other hand, the p75NTR ectodomain is neuroprotective against amyloid-beta toxicity in AD (Yao et al., 2015). Similarly, future CNTNAP2-ecto-based therapies could prevent seizures, modulate intracellular Ca^2+^, and activate PMCAs in PMCA-associated disorders.

### Ca^2+^ extrusion and neuronal synchrony in neurodevelopmental disorders

We show that PMCAs are novel binding partners for CNTNAP2-ecto. PMCAs have been associated with autism, ataxia, deafness, and cardiovascular disease. Specifically, PMCA2 has been implicated in ASD (Takata et al., 2018). We found a reduction in PMCA2 levels in the *Cntnap2* KO mice cortex at P28. Interestingly, a reduction in the protein levels of PMCA2 has been observed in postmortem brain tissue from ASD patients (Voineagu et al., 2011). Moreover, decreased gene expression of *ATP2B2* has also been observed in lymphoblasts of autism patients (Stafford et al., 2017).

We show that CNTNAP2-ecto modulates neuronal network synchrony. Alterations in neuronal synchrony are thought to occur in ASD, SZ, (Mathalon and Sohal, 2015) and epileptic seizures (McCormick and Contreras, 2001), disorders associated with CNTNAP2 genetic variation. Abnormal synchrony has been reported in multiple mouse models of neurodevelopmental disorders, including *Cntnap2* KO (Penagarikano et al., 2011), *Fmrp* (Goncalves et al., 2013), 22q11 deletion (Sigurdsson et al., 2010), and neuroligin 3 R451C (Gutierrez et al., 2009), suggesting that abnormal synchrony is a common pathophysiological mechanism in neurodevelopmental disorders.

As such, abnormal levels of CNTNAP2-ecto, either caused by anomalous CNTNAP2 expression or shedding, could lead to altered synchrony, or alter seizure susceptibility. Human subjects with *CNTNAP2* risk variants showed reduced long-range and increased local connectivity in cortical circuits (Scott-Van Zeeland et al., 2010), while healthy individuals with the risk allele had increased activation of specific cortical regions (Whalley et al., 2011).

## Supporting information

SuppVideo1

SuppVideo2

## ACKNOWLEDGMENTS

This work was supported by grant MH097216 from the NIH-NIMH (to P.P.), by an individual Biomedical Research Award from the Martwell Foundation (to J.N.S), by NIH Grant R01 MH092906 (to D.C.), and by the NJ Commission on Traumatic Brain Injury Research (CBIR16PIL035), the Fred & Santa Barile Children’s Medical Research Trust (to D. C.), and the Robert Wood Johnson Foundation (Grant 74260) in support of the Child Health Institute of New Jersey. SIM imaging work was performed at the Northwestern University Center for Advanced Microscopy, generously supported by NCI CCSG P30 CA060553. The Nikon N-SIM system was purchased through the support of NIH 1S10OD016342-01. We thank Drs. Joshua Zachary Rappoport and Constadina Arvanitis for help with SIM imaging and live Ca^2+^ imaging.

## AUTHOR CONTRIBUTIONS

MDMS, MdS, OV, BPS, RG, MF, KM, NK, EAH, ASC, DC conducted experiments; PP, MDMS, OV, JNS, designed experiments; MDMS, MdS, PP, SFL, JNS analyzed the data; PP and MDMS wrote the paper.

## MATERIALS AND METHODS

### CONTACT FOR REAGENT AND RESOURCE SHARING

Further information and requests for resources and reagents should be directed to and will be fulfilled by the Lead Contact, Peter Penzes.

### EXPERIMENTAL MODEL AND SUBJECT DETAILS

#### Cntnap2 knockout mice

*Cntnap2* KO mice were generated and kindly donated by Elior Peles (Weizmann Institute of Science, Israel). The colony was genotyped following the primers and conditions previously described (Poliak et al., 2003), and it was maintained in their original strain, CD1.

#### Neuronal culture

High density (100,000 cells/cm^2^) cortical neuron cultures were prepared from P0 *Cntnap2* WT or KO mice or Sprague-Dawley rat E18 embryos as described previously (Srivastava et al., 2011). Briefly, neurons were plated onto coverslips coated with poly-D-lysine (0.2 mg/mL; Sigma), in feeding medium (Neurobasal medium supplemented with B27 (Invitrogen), 0.5 mM glutamine and penicillin/streptomycin). 200 μM DL-APV (Abcam) was added to the medium 4 days later. Half of the feeding medium+APV was replaced twice a week until the neurons were used.

#### Neuronal transfections

Cortical neurons were transfected at DIV 21 using Lipofectamine 2000 following the manufacturer’s recommendations, and the neurons were maintained in the feeding media for 2 days after the transfection. Neurons were then fixed in 4% formaldehyde + 4% sucrose in PBS for 10 min and washed 3 times with PBS for 10 min. Coverslips were then processed for immunostaining. Only neurons that exhibited a pyramidal asymmetric morphology, with a single long apical and highly branching protrusion, and many basal shorter protrusions radiating from the soma, were selected for further analysis. Any signs of poor neuronal health excluded the cell from quantification. Rats and mice were used in accordance with IACUC and national guidelines and regulations under approved protocols.

### METHOD DETAILS

#### Antibodies, plasmids and CSF samples

The following antibodies were purchased: CNTNAP2 (C-terminal, rabbit polyclonal, Millipore, Darmstadt, Germany), CNTNAP2 (N-terminal, mouse monoclonal, Neuromab, Davis, California), actin (mouse monoclonal, Abcam, Cambridge, United Kingdom), GFP (chicken polyclonal, ab13972; Abcam), Flag (Mouse, Sigma), vGLUT1 (Guinea Pig, Synaptic System Antibodies), PMCA2 (Rabbit, Thermo Fisher), CNTN-1 (Abcam). Plasmids used in this study were: pEGFP-N1 (Clontech, Mountain View, California); Flag-CNTNAP2 (generated by subcloning human CNTNAP2 cDNA from a plasmid kindly provided by Dr. Elior Peles, Weizmann Institute of Science, Israel) (Gao et al., 2018); PMCA2w/b and GCamP6s (both from Addgene); and DsRedExpress2 (Clontech). Pooled murine (CD1) and remnants of human CSF were obtained from Bioreclamation IVT.

#### Biological process enrichment analysis

The gene annotation enrichment analysis tool DAVID v6.8 was used to perform biological process GO term enrichment (Huang da et al., 2009). For the secretome analysis, the list of detected proteins previously published in (Kuhn et al., 2016) was used as an initial dataset. For the hCSF analysis, a sample from a healthy individual was obtained from BIoreclamation IVT, and MS was performed. For comparison, a published hCSF proteome (Kroksveen et al., 2017) was included in Fig.S1A-C with a list of 2071 IDs. In order to avoid any nomenclature discrepancies, all IDs were translated into *entrez* gene ID, thus we used *H. sapiens* as the background list except for the membrane-anchored analysis, for which we used as background a list of transmembrane and GPI-anchored proteins obtained from UniProt. The –log_10_ of the Benjamini corrected *P*-value was calculated and graphed for the top 5 non-redundant GOTERM_BP_FAT categories, where statistical significance is reached at a value of 1.3.

#### Disease enrichment analysis

A hypergeometric test was used to calculate the probability of finding an over- or under-enrichment of disease-relevant genes in the secretome, hCSF, and the published hCSF gene ID lists and sub-lists (including total, membrane-anchored, and secreted proteins). As disease-relevant gene lists, we used all *de novo* gene lists from the supplementary data compiled in (Genovese et al., 2016). Only *de novo* variants affecting protein-coding regions or splice sites were included (that is, missense, frameshift, and splice site mutations, altered stop codons, altered start codons, insertions, and deletions).

#### Protein-protein interaction network analysis with STRING

STRING v10.5 was used to perform protein-protein interaction (PPI) analysis. The confidence of the PPI is shown by the network edges, using “experiments” and “databases” as the active interaction sources, with medium confidence. We adjusted the network to fit 4 clusters and used the k means clustering method for creating the PPI network to provide a global and intuitive understanding of the functional properties of the proteins belonging to the synaptic sheddome, defined as the overlap between the hCSF dataset (1416 IDs) and the PSD proteome (Bayes et al., 2012).

#### Chemical LTP induction protocol

High density, 24 days *in vitro* (DIV) rat cortical neurons were accommodated for 30 min in aCSF (containing in mM: NaCl 125, KCl 2.5, CaCl_2_ 2, MgCl_2_ 1.25, glucose 11, NaHCO_3_ 26.2, NaH_2_PO_4_ 1, Hepes 10, 200 μM APV, pH 7.4). Then, LTP was induced chemically by the application of a chemical cocktail containing 100 μM picrotoxin, 1 µM strychnine,10 μM glycine, 0 Mg^2+^, 0 APV, in aCSF) for 30 min. After this, extracellular media proteins were collected and concentrated by the TCA method and cells were lysed in RIPA buffer. When inhibitors or antagonists were employed, they were added in the accommodation to aCSF phase as well as during the cLTP phase. The following chemicals were used: dynasore (Sigma, 100 μM), APV (Sigma, 200 μM), NBQX (Tocris, 10 μM), TAPI-0 (Enzo, 25 μM, which inhibits MMP13, MMP9, MMP1, TACE and MMP3 with decreasing affinities K_i_ in nM 0.2, 0.5, 6, 8.8, 68 respectively), GM6001 (Calbiochem, 50 μM), MMP9 inhibitor I (Calbiochem,10 μM).

#### Precipitation of extracellular media proteins by TCA method

Supernatants were centrifuged for 3 min at 16,000 rpm to eliminate suspended cells. 0.015% deoxycholate was incubated with the supernatants for 5 min. TCA was added to the supernatants to a final concentration of 20%. After an incubation period of 1 h, the samples were centrifuged at 18,000×g. The pellets were washed three times with ice cold acetone. Remaining acetone was evaporated, and the pellets were dissolved in sample buffer.

#### Separation of membrane and soluble fractions from tissue

The protocol described by Suzuki *et al*. (Suzuki et al., 2012) was used with slight modifications. One-month old male WT and *Cntnap2* KO mice were dissected, and cortices were homogenized with a dounce homogenizer in a Tris-based buffer (50 mM Tris-HCl, pH 7.6, 150 mM NaCl, and Roche protease inhibitor cocktail). They were centrifuged at 1500 g for 10 min, and the supernatants were subsequently centrifuged at 100,000×g for 1 h. The pellet containing the membranes and the supernatants containing soluble proteins were analyzed by western blotting.

#### Immunocytochemistry

24 DIV neurons were first washed in PBS and then fixed in 4% formaldehyde-4% sucrose-PBS for 10 min. Fixed neurons were then permeabilized and blocked simultaneously (0.1% BSA, 4% NGS, 0.1% Triton-X-100 in PBS, 1 h at RT), followed by incubation of primary antibodies overnight at 4 °C. The coverslips were washed 3 times with PBS and incubated with the corresponding fluorophore-secondary antibodies (45 min, RT) (Alexa Fluor^®^488, Alexa Fluor^®^596, LifeTechnologies, Carlsbad, California), as described previously. The coverslips were then mounted onto slides using ProLong Antifade reagent (Invitrogen), and stored at 4 °C until the acquisition of the images. All images were acquired in the linear range of fluorescence intensity. When vGlut1 antibody was used, incubation with primary antibodies and the corresponding secondary antibodies was performed in two consecutive days to avoid cross-reaction of chicken Alexa Fluor^®^488 with guinea pig vGlut1 primary antibody.

#### SIM imaging and analysis

3-week old *Cntnap2* KO cortical neurons were transfected with GFP and an extracellularly tagged Flag-CNTNAP2. Then, after a 30-min accommodation of the cells in aCSF (containing in mM: NaCl 125, KCl 2.5, CaCl_2_ 2, MgCl_2_ 1.25, glucose 11, NaHCO_3_ 26.2, NaH_2_PO_4_ 1, Hepes 10, pH 7.4), LTP was induced chemically (100 µM picrotoxin, 1 µM strychnine,10 µM glycine, 0 Mg^2+^, 0 APV in aCSF) for 30 min. After this, cells were fixed and stained for GFP, vGLUT1, Flag, and CNTNAP2 C-terminal antibody. Images were taken with an N-SIM microscope at the Nikon Imaging Center at Northwestern University, as previously described (Smith et al., 2014). Acquisition was set to 10 MHz, 14 bit with EM gain, and no binning. Autoexposure was kept at 100-300 ms, and the EM gain multiplier restrained below 300. Laser power was adjusted to keep look up tables (LUTs) within the first quarter of the scale. Reconstruction parameters Illumination Modulation Contrast, High Resolution Noise Suppression, and Out of Focus Blur Suppression (0.96, 1.19, and 0.17, respectively) were kept consistent across experiments and imaging sessions, as described (Smith et al., 2014).

Images were analyzed with ImageJ. Line scans were used to generate fluorescence intensity profile plots to assess colocalization. CNTNAP2-ecto was identified using the AND, OR, and XOR functions, in order to identify signal that contained only the flag staining (N-terminal or extracellular) and no C-terminal/intracellular signal. This flag signal void of CNTNAP2 C-terminal signal was considered to correspond to CNTNAP2-ecto and was analyzed in two different compartments: GFP or overlap between GFP and vGLUT1 (synaptic). The area occupied by CNTNAP2-ecto and the number of punctae were measured within the GFP and synaptic regions.

#### Fractionation protocol for neuronal culture

We followed a protocol described elsewhere, with slight modifications (Won et al., 2016). Briefly, 3-week old rat cortical neurons were rinsed in PBS and cells were collected in a PBS solution containing 1 mM MgCl_2_ and 0.1 mM CaCl_2_, protease inhibitor cocktail, and phosphatase inhibitor cocktail. Cells were harvested after centrifugation at 16,000×g for 2 min, and lysed for 15 min in hypotonic buffer (10 mM Tris, 1 mM MgCl_2_ and 0.1 CaCl_2,_ protease inhibitor cocktail, and phosphatase inhibitor cocktail, pH 7.5). Membranes were pelleted by centrifugation (20,000×g, 20 min) and lysed for 15 min in lysis buffer (PBS, 1 mM MgCl_2_, 0.1 CaCl_2,_ 1% Triton-X-100, protease inhibitor cocktail, and phosphatase inhibitor cocktail). Then, samples were centrifuged at 20,000×g for 20 min, finding the extrasynaptic fraction in the supernatant, and the insoluble pellet containing the synaptic fraction was dissolved in PBS, 1 mM MgCl_2_, 0.1 CaCl_2,_ 1% Triton X-100, and 1% SDS. Whole cell lysates and extrasynaptic and synaptic fractions were analyzed by western blotting.

#### CNTNAP2-ecto protein production

HEK cells were plated in PDL-coated 10 cm^2^ dishes and transfected with Flag-CNTNAP2 using lipofectamine following the manufacturer’s recommendations. After a 2-day transfection, supernatants from CNTNAP2-transfected and non-transfected cultures were collected, centrifuged at 700×g, and filtered through a 0.22 µm filter. In order to concentrate the supernatants to a smaller and more manageable volume, samples were spin-filtered with molecular cut-off filters of 100 kD (Millipore). The concentrated supernatant was then affinity-purified with flag-agarose beads (Sigma) following the manufacturer’s instructions. Briefly, beads were washed and rinsed, added to the concentrated supernatants and rocked at 4 °C for 4 h. After a 2 min centrifugation at 2,000×g, the beads were rinsed 3 times with TBS and finally the proteins were eluted with 100 mM Gly, pH 3.5, and beads were removed by centrifugation. 1 M Tris, pH 8.5, was added to the eluate to neutralize the solution. Finally, the samples containing the purified CNTNAP2-ecto and the vehicle control were dialyzed 3 times using 100 kDa spin filters (Millipore) to remove the glycine and substitute it with a TBS solution.

#### Expression and purification of CNTNAP2-ecto

The construct encoding the entire extracellular domain of CNTNAP2 from residue 28 to 1261 was fused to the Fc portion of human IgG1 as detailed elsewhere (Rubio-Marrero et al., 2016). This construct was transfected into HEK293GnTI-cells and the cells were selected by growth in the antibiotic G418. For large scale protein expression, stable cells were maintained at 37 °C and 5% (v/v) CO_2_ in Dulbecco’s modified Eagle’s medium containing up to 5% (v/v) FBS. CNTNAP2-Fc secreted in the medium was affinity purified using ProteinA-CaptivA TM PriMAB (RepliGen, Waltham, MA) and subsequently cleaved with recombinant 3C protease to remove the Fc fragment and leave the mature (i.e., no leader peptide) extracellular domain of CNTNAP2. The purified protein was buffer exchanged into 10 mM Hepes, pH 7.4, and 150 mM NaCl, concentrated to ∼3 to ∼6 mg/mL, and stored at 4 °C until used.

#### Affinity chromatography and mass spectrometry

##### Tissue preparation

Cortices from 20 6-8-week old male C57BL6J mice were homogenized on ice with a dounce homogenizer in buffer HB (20 mM Hepes, pH 7.4, 320 mM sucrose, 5 mM EDTA, protease inhibitors). Homogenates were centrifuged at 3,000×g for 15 min at 4 °C to remove nuclei and cell debris. Supernatant was further centrifuged at 38,400×g (15 min, 4 °C) and the pellet was resuspended in buffer SB1 (20 mM Hepes, pH 7.4, 1 M KI, 5 mM EDTA, protease inhibitors) to remove peripheral proteins. This homogenate was then centrifuged at 38,400×g (15 min, 4 °C) to collect membranes. Pellet containing membranes was washed with buffer SB2 (20 mM Hepes, pH 7.4, 5 mM EDTA, protease inhibitors) to remove KI and it was centrifuged again at 38,400×g for 30 min at 4 °C. The pellet was then resuspended in buffer RB (20 mM Hepes, pH 7.4, 100 mM NaCl, 5 mM EDTA, 5 mM EGTA, 1% CHAPS, protease inhibitors) and rotated for 1.5 h at 4 °C to solubilize the membranes. Finally, the homogenate was ultracentrifuged at 100,000×g for 1 h at 4 °C to remove insoluble materials. The final supernatant was split in halves. Vehicle or CNTNAP2-ecto was added to the membrane homogenate and incubated overnight at 4 °C. The following morning, the same amount of flag-agarose beads (Sigma) was added to each sample and incubated with rotation for 1.5 h at 4 °C. After capture, samples were centrifuged at 2,000×g for 1 min, and the beads were rinsed with buffer RB 3 times. The samples containing the beads were passed through disposable chromatography columns (BioRad) and then bait and prey proteins were eluted by adding a solution containing 100 mM glycine, 1% CHAPS, pH 2.5, 3 times. The eluate was precipitated by adding trichloracetic acid to a final concentration of 20% over 1 h at 4 °C, and the solution was then centrifuged at 18,000×g for 1 h at 4 °C. Finally, the pellet was washed 3 times with ice cold acetone, and the samples were kept at RT to dry and stored at −80 °C until they were analyzed by mass spectrometry.

##### Sample preparation for LC-MS/MS analysis

We followed the protocol described elsewhere (Hickox et al., 2017). The precipitated protein pellets were solubilized in 100 μL of 8 M urea for 30 min; 100 μL of 0.2% ProteaseMAX (Promega) was then added, and the mixture was incubated for an additional 2 h. The protein extracts were reduced and alkylated as described previously (Chen et al., 2008), followed by the addition of 300 μL of 50 mM ammonium bicarbonate, 5 μL of 1% ProteaseMAX, and 2 μg of sequence-grade trypsin (Promega). Samples were digested overnight in a 37 °C thermomixer (Eppendorf). For Orbitrap Fusion Tribrid MS analysis, the tryptic peptides were purified with Pierce C18 spin columns and fractionated with increasing acetonitrile (ACN) concentrations (30% and 70%). Up to 3 μg of each fraction was auto-sampler loaded with a Thermo Fisher EASY nLC 1000 or nLC 1200 UPLC pump onto a vented Acclaim Pepmap 100, 75 μm by 2 cm, nanoViper trap column coupled to a nanoViper analytical column (Thermo Fisher 164570, 3 μm, 100 Å, C18, 0.075 mm, 500 mm) with stainless steel emitter tip assembled on the Nanospray Flex Ion Source with a spray voltage of 2000 V. Buffer A contained 94.785% H_2_O with 5% ACN and 0.125% formic acid (FA), and buffer B contained 99.875% ACN with 0.125% FA. The chromatographic run was 4 h in total with the following profile: 0–7% for 7 min, 10% for 6 min, 25% for 160 min, 33% for 40 min, 50% for 7 min, 95% for 5 min, and 95% again for 15 min. Additional MS parameters include: ion transfer tube temp=300 °C, Easy-IC internal mass calibration, default charge state=2, and cycle time=3 s. Detector type set to Orbitrap, with 60 K resolution, with wide quad isolation, mass range=normal, scan range=300–1500 m/z, max injection time=50 ms, AGC target=200,000, microscans=1, S-lens RF level=60, without source fragmentation, and datatype=positive and centroid. MIPS was set to ‘on’, and included charge states=2–6 (reject unassigned). Dynamic exclusion was enabled, with n=1 for 30 and 45 s exclusion duration at 10 ppm for high and low, respectively. Precursor selection decision=most intense, top 20, isolation window=1.6, scan range=auto normal, first mass=110, collision energy 30%, CID, Detector type=ion trap, Orbitrap resolution=30K, IT scan rate=rapid, max injection time=75 ms, AGCtarget=10,000, Q=0.25, inject ions for all available parallelizable time.

##### Tandem mass spectra analysis

Peptide spectral files from pooled samples or from biological replicates were combined for database searching. Raw spectrum files were extracted into MS1 and MS2 files using in-house program RawXtractor or RawConverter (http://fields.scripps.edu/downloads.php) (He et al., 2015) and the tandem mass spectra were searched against UniProt’s mouse protein database (downloaded on 09– 21-2015; UniProt Consortium, 2015) and matched to sequences using the ProLuCID/SEQUEST algorithm (ProLuCID version 3.1; (Eng et al., 1994) with 50 ppm peptide mass tolerance for precursor ions and 600 ppm for fragment ions. The human CNTNAP2 amino acid sequence was manually added to the database:

MQAAPRAGCGAALLLWIVSSCLCRAWTAPSTSQKCDEPLVSGLPHVAFSSSSSISGSYSPGYAKINKRGGAGGWSPSDSDHYQWLQVDFGNRKQISAIATQGRYSSSDWVTQYRMLYSDTGRNWKPYHQDGNIWAFPGNINSDGVVRHELQHPIIARYVRIVPLDWNGEGRIGLRIEVYGCSYWADVINFDGHVVLPYRFRNKKMKTLKDVIALNFKTSESEGVILHGEGQQGDYITLELKKAKLVLSLNLGSNQLGPIYGHTSVMTGSLLDDHHWHSVVIERQGRSINLTLDRSMQHFRTNGEFDYLDLDYEITFGGIPFSGKPSSSSRKNFKGCMESINYNGVNITDLARRKKLEPSNVGNLSFSCVEPYTVPVFFNATSYLEVPGRLNQDLFSVSFQFRTWNPNGLLVFSHFADNLGNVEIDLTESKVGVHINITQTKMSQIDISSGSGLNDGQWHEVRFLAKENFAILTIDGDEASAVRTNSPLQVKTGEKYFFGGFLNQMNNSSHSVLQPSFQGCMQLIQVDDQLVNLYEVAQRKPGSFANVSIDMCAIIDRCVPNHCEHGGKCSQTWDSFKCTCDETGYSGATCHNSIYEPSCEAYKHLGQTSNYYWIDPDGSGPLGPLKVYCNMTEDKVWTIVSHDLQMQTPVVGYNPEKYSVTQLVYSASMDQISAITDSAEYCEQYVSYFCKMSRLLNTPDGSPYTWWVGKANEKHYYWGGSGPGIQKCACGIERNCTDPKYYCNCDADYKQWRKDAGFLSYKDHLPVSQVVVGDTDRQGSEAKLSVGPLRCQGDRNYWNAASFPNPSSYLHF STFQGETSADISFYFKTLTPWGVFLENMGKEDFIKLELKSATEVSFSFDVGNGPVEIVVRSPTPLNDDQWHRVTAERNVKQASLQVDRLPQQIRKAPTEGHTRLELYSQLFVGGAGGQQGFLGCIRSLRMNGVTLDLEERAKVTSGFISGCSGHCTSYGTNCENGGKCLERYHGYSCDCSNTAYDGTFCNKDVGAFFEEGMWLRYNFQAPATNARDSSSRVDNAPDQQNSHPDLAQEEIRFSFSTTKAPCILLYISSFTTDFLAVLVKPTGSLQIRYNLGGTREPYNIDVDHRNMANGQPHSVNITRHEKTIFLKLDHYPSVSYHLPSSSDTLFNSPKSLFLGKVIETGKIDQEIHKYNTPGFTGCLSRVQFNQIAPLKAALRQTNASAHVHIQGELVESNCGASPLTLSPMSSATDPWHLDHLDSASADFPYNPGQGQAIRNGVNRNSAIIGGVIAVVIFTILCTLV FLIRYMFRHK GTYHTNEAKGAESAESADAA IMNNDPNFTETIDESKKEWLI

The search space included all fully and half-tryptic or non-tryptic (for CNTNAP2 sequence mapping) peptide candidates that fell within the mass tolerance window with no miscleavage constraint. Peptides were assembled and filtered with DTASelect2 (version 2.1.3) (Cociorva et al., 2007; Tabb et al., 2002) through Integrated Proteomics Pipeline (IP2 version 3, Integrated Proteomics Applications, http://www.integratedproteomics.com). To estimate peptide probabilities and false discovery rates (FDR) accurately, we used a target/decoy database containing the reversed sequences of all the proteins appended to the target database (Peng et al., 2003). Each protein identified was required to have a minimum of one peptide of minimal length of 6 amino acid residues; however, this peptide had to be an excellent match with a FDR <0.001 and at least one excellent peptide match. After the peptide/spectrum matches were filtered, we estimated that the protein FDRs were ≤1% for each sample analysis. Resulting protein lists included subset proteins to allow for consideration of all possible protein forms implicated by a given peptide identified from the complex protein mixtures.

#### Immunoprecipitation in HEK293 cells

One 10 cm^2^ dish was seeded with HEK293T cells. At 70% confluency, cells were transfected with PMCA2w/b with lipofectamine 2000 following the manufacturer’s instructions. After a 2-day transfection, cells were lysed in Hepes buffer (20 mM Hepes, 1% CHAPS, 100 mM NaCl, 5 mM EDTA, protease inhibitor cocktail (Roche), phosphatase inhibitor cocktail III (Sigma)). The lysate was precleared for 30 min at 4 °C with anti-flag M2 affinity beads (Sigma). CNTNAP2-ecto or vehicle was incubated with the cell lysate overnight. Then, flag beads were added to the lysate and incubated for 2 h. The beads were washed 3 times with Hepes buffer, and finally, sample buffer was added to the beads, and the mixture was heated for 20 min at 40 °C to elute the proteins. Proteins were detected by western blotting.

#### Cytosolic Ca^2+^ imaging in HEK293 cells

One day after seeding, cells were transfected with GFP-PMCA2w/b. After 2 days, cells were loaded with 1 µM Fura-2AM, a fluorescent Ca^2+^ indicator used to measure cytosolic free Ca^2+^ levels. Cells were rinsed twice with recording solution containing, in mM, NaCl 145, KCl 5, CaCl_2_ 2, MgCl_2_ 1, glucose 10, Hepes 10, pH 7.4. The glass-bottomed dish containing the cells was placed on the stage of a C2 Nikon microscope where the temperature was maintained at 37 °C. Images were acquired using a 20X objective. After 60 s of basal recording, the cells were treated with either vehicle or CNTNAP2-pNTF, after which 0.5 mM ATP (Sigma) was added to elicit a cytosolic Ca^2+^ signal. Acquired images were analyzed with NIS elements (Nikon), selecting both PMCA2 transfected and non-transfected cells.

#### Cytosolic Ca^2+^ imaging in dissociated rat cortical neurons

At day-in-culture 17, rat cortical neurons were transfected with the cytosolic genetic Ca^2+^ indicator GCaMP6s and the red cell fill DsRed Express2. After 2 days, cells were cells were rinsed twice with aCSF containing (in mM) NaCl 125, KCl 2.5, CaCl_2_ 2, MgCl_2_ 1.25, glucose 11, NaHCO_3_ 26.2, NaH_2_PO_4_ 1, Hepes 10, pH 7.4. The glass-bottomed dish containing the neurons was placed on the stage of a microscope where the temperature was maintained at 37 °C. After 60 s of basal recording, the cells were treated with either vehicle or CNTNAP2-pNTF, after which 30 mM KCl was added to elicit a cytosolic Ca^2+^ signal. Acquired images were analyzed with NIS elements (Nikon). For the analysis of Ca^2+^ in soma, a C2 Nikon microscope and a 20X objective were used.

#### Brain slice preparation

Animals were deeply anesthetized with 5% isoflurane in O_2_ and decapitated. The brain was rapidly extracted and placed on ice-cold modified aCSF containing (in mM) NaCl 125, NaHCO_3_ 26, glucose 11, KCl 2.5, NaH_2_PO_4_ 1.25, CaCl_2_ 2.5, MgCl_2_ 6, and kynurenic acid 1 for 2-3 min and then glued on a vibratome VT1000S (Leica). The slicing chamber was filled with the same ice-cold aCSF. Slices then recovered at 34 **°**C for at least 45 min in classic aCSF (in mM, NaCl 125, NaHCO_3_ 26, glucose 11, KCl 2.5, NaH_2_PO_4_ 1.25, CaCl_2_ 2.5, MgCl_2_ 1.3) before incubation with the Ca^2+^ sensor Cal520-AM (15 mM, AAT Bioquest) and the glial indicator SR-101 (1 μM, Sigma) for 45 min at 34 **°**C with constant bubbling of 95% O_2_/5% CO_2_ in a custom made incubator. Slices were then pre-incubated for 1 h with 10 nM of the CNTNAP2-ecto or vehicle at 34 **°**C before imaging. Slices were transferred to an imaging chamber (Warner) with a constant flow of 3 mL/min of 95% O_2_/5% CO_2_ in aCSF under a Nikon A1R multiphoton microscope, immersion objective 25X, NA 1.1. A chameleon laser was tuned at 820 nm and the power was set around 12-16 mW. aCSF was kept at 37.5 **°**C by a heated controller (TC344, Warner Instrument) to have a final temperature of 32-33 **°**C in the imaged sample. Slices were allowed to stabilize for 5 min in the chamber before recording. The 30 mM KCl evoked responses were acquired at 2 frames/s, and spontaneous activity was recorded at a speed of 3 frames/s with 7.5 mM of KCl to induce spontaneous network events (Feldt Muldoon et al., 2013).

#### Analysis of spontaneous activity and synchrony

Regions of interest were traced manually on somas based on the maximum projection obtained in ImageJ, blinded for the experimental conditions. Somas stained for the red glial dye SR-101 were not included in the analysis. The background was subtracted, and the mean intensity was used as the intensity of the fluorescence for each time point. The normalized data were analyzed in Matlab (Matwork) using a homemade routine based on the “findpeak” functions of the software. The minimum variation of intensity was set at 6% for neuronal cultures and at 10% for slices. ROIs presenting no transients during the time course of the movie were discarded from the analysis. To prevent bias while detecting the time windows when the neuronal network was synchronized, we used a method based on 10,000 random permutations over time of the binary Ca^2+^ transients for each neuron. The generated distribution of the number of events per frame is used to determine a threshold of significant synchrony corresponding to *Threshold Synchrony = µ + 3 σ* where μ is the average number of events per frame in the 10,000 shuffled data, and σ is the standard deviation of number of events per frame in the shuffled data. The synchrony ratio corresponds to the fraction of transients during the synchrony phases divided by the total number of events (Fig.S6).

### QUANTIFICATION AND STATISTICAL ANALYSIS

Statistical analysis was performed using GraphPad Prism 5.0 (La Jolla, USA). To compare differences between two groups, Student’s *t*-test (Gaussian distribution of data) or a Mann Whitney test (non-Gaussian distribution of data) was used. Data distribution was considered Gaussian after passing the Kolmogorov-Smirnov test or the F-test to compare variances when n<5. To compare multiple variables, two-way ANOVA was used, followed by a Newman-Keuls *post hoc* test to compare replicate means unless otherwise specified. Statistical significance was defined as *P*<0.05.

### KEY RESOURCES TABLE

To be provided upon acceptance.

## SUPPLEMENTAL FIGURE LEGENDS

**Figure S1, Related to Figure 1. A.** Venn diagram showing that, of the 2072 proteins taken from a published human CSF dataset, 860 could potentially undergo ectodomain shedding. GO analysis performed for originally membrane-anchored proteins shows cell adhesion and axon guidance as the top enriched categories. **B.** The membrane-anchored fraction of the published hCSF shows an enrichment for ASD-related genes, but no significant enrichment is observed in the soluble or secreted proteins. **C.** Hypergeometric test shows a highly significant overlap between the neuronal sheddome *in vitro* and the published hCSF sheddome (139 proteins). **D.** GO analysis performed for soluble proteins in the neuronal secretome, the hCSF and the published hCSF shows extracellular matrix organization and platelet degranulation as the top enriched categories. **E.** Hypergeometric tests run in the three datasets analyzed (neuronal secretome, hCSF, and previously published hCSF) show that only the membrane-anchored proteome is enriched for ASD-related risk factors taken from the SFARI database (category 1-5). **F.** The number of proteins per biological process (Y axis) in the neuronal, hCSF, and previously published hCSF sheddomes shows that cell adhesion is the process with the highest number of proteins in the three datasets.

**Figure S2. A.** Cell lysates and extracellular media extracted from HEK293 cells transfected with WT and ΔCT CNTNAP2 and the CDFE ASD associated mutation 1253* after a two-day transfection. Samples were immunoblotted for CNTNAP2 N-term antibody and CNTNAP2 C-term antibody. * used to mark the intracellular C-terminal fragment produced after ectodomain shedding. # used to mark the CNTNAP2-ecto found in the WT, dCT and the 1253* samples. Extracellular media samples were devoid of actin signal, indicating no cell death or debri present in the samples. Moreover, there was no signal with the C-terminal antibody in the media samples, confirming that the signal observed with the N-term antibody corresponds to the CNTNAP2-ecto and not to the full length. **B.** N2a cells transfected with CNTNAP2 display CNTNAP2 ectodomain shedding that is blocked with the inhibitors GM6001, TAPI-O and MMP9 inhibitor I, corroborating our data obtained in cultured neurons.

**Figure S3, Related to Figure 3. A.** Analysis of the localization of CNTNAP2-ecto shows its presence both within the cell as the full-length CNTNAP2 (45%) and in the extracellular space adjacent to the neuron (55%), demonstrating CNTNAP2 ectodomain shedding by SIM, a superresolution imaging technique. **B.** Analysis of the localization of CNTNAP2-ecto by SIM imaging shows that it is localized extracellularly, both well outside the axonal compartment and close to synaptic boutons, and also within axons.

**Figure S4, Related to Figure 4. A.** Pull down experiment of HEK293 cells transfected with PMCA2, incubated with WT CNTNAP2-ecto or disease associated CNTNAP2 1253* demonstrates that both forms of CNTNAP2 interact with PMCA2. **B.** Confocal images of dissociated rat cortical neurons shows colocalization of PMCA2 and CNTNAP2. **C.** PMCA2 expression levels in *Cntnap2* WT and KO mice. WT mice show increasing amounts of PMCA2 with age, while KO mice present decreased levels of PMCA2 at age P28 compared with the WT counterparts. Data are represented as mean±SEM, n=3, *t*-test \**P*=0.0349. **D.** Allen brain atlas images of CNTNAP2 and PMCA2 show a similar pattern of expression of both proteins in 8^th^ sagittal slice of P56 mice brains. **E.** Experimental protocol employed for Ca^2+^ imaging in HEK293 cells transfected with PMCA2 and treated with vehicle or CNTNAP2-ecto. Kinetic parameters such as peak, t_max_, and area under the curve (AUC) show no difference between vehicle and CNTNAP2-ecto-treated PMCA2-transfected HEK293 cells. Data are represented as mean±SEM. <u>Peak data:</u> N=7 cultures, one-way ANOVA \**P*<0.0001, Newman-Keuls *post-hoc* test, ns. <u>t_max_:</u> N=7 cultures, Kruskal-Wallis test \**P*<0.02, Dunn’s *post-hoc* test, ns. <u>AUC:</u> N=6 cultures, one-way ANOVA \**P*=0.3149, Newman-Keuls *post-hoc* test, ns.

**Figure S5, Related to Figure 5. A.** Scheme of the protocol followed for Ca^2+^ imaging in dissociated rat cortical neurons. Cells were transfected with the Ca^2+^ indicator GCaMP6s and the cell fill DsRedExpress2. After two days, they were incubated with vehicle (Veh) or CNTNAP2-ecto (Ecto) for 5 min. KCl at 30 mM was then added to the cells to depolarize neurons and elicit a Ca^2+^ peak. **B.** Concentration response curve of CNTNAP2-ecto at 3, 10, and 30 nM indicates that 10 nM is the most efficient concentration to decrease the t_1/2_ off; consequently, the rest of the experiments were performed at this concentration. Data are represented as mean±SEM, n=25-47 neurons, one-way ANOVA *P*=0.0312, Newman-Keuls *post-hoc* test \**P*=0.0312. **C.** Other kinetic parameters analyzed at CNTNAP2-ecto 10 nM, such as the peak and the t_max_, were not altered by the incubation with CNTNAP-ecto, confirming that the effect of the CNTNAP2-ecto is specific to the Ca^2+^ extrusion phase. Data are represented as mean±SEM, n=39/23 neurons, peak: Mann-Whitney test, *P*=0.1370; t_max_ Mann-Whitney test, *P*=0.2417. **D.** Using the cell fill DsRedExpress2 to dissociate the effect of CNTNAP2-ecto in pyramidal *vs* inhibitory neurons based on their different morphology, we found that the effect exerted by the CNTNAP2-ecto on the t_1/2_ off is approximately the same (23.7% *vs* 22.5% reduction), but only significant in pyramidal neurons, probably due to the variability encountered in inhibitory neurons. Data are represented as mean±SEM. Pyramidal neuron analysis: Mann-Whitney test, n=28-23, \**P*=0.0285. Inhibitory neuron analysis: *t*-test, n=11-9, *P*=0.1781.

**Figure S6. A.** Representative heat map of the co-activation between pairs of cells observed by spontaneous Ca^2+^ imaging on control primary cortical neurons. **B.** Co-activation map after a random shuffling in time of the Ca^2+^ transients. **C.** Raster plot of the real Ca^2+^ transients and **D.** after reshuffling of the peaks. **E.** After 10,000 reshufflings of Ca^2+^ transients, a distribution of the random surrogate data is used to determine an unbiased threshold of significant synchrony corresponding to the average of the random distribution plus 3 times the standard deviation.

