## Supplementary figures and images for "CNTNAP2 ectodomain, detected in neuronal and CSF sheddomes, modulates Ca^2+^ dynamics and network synchrony"

### Fig.S1

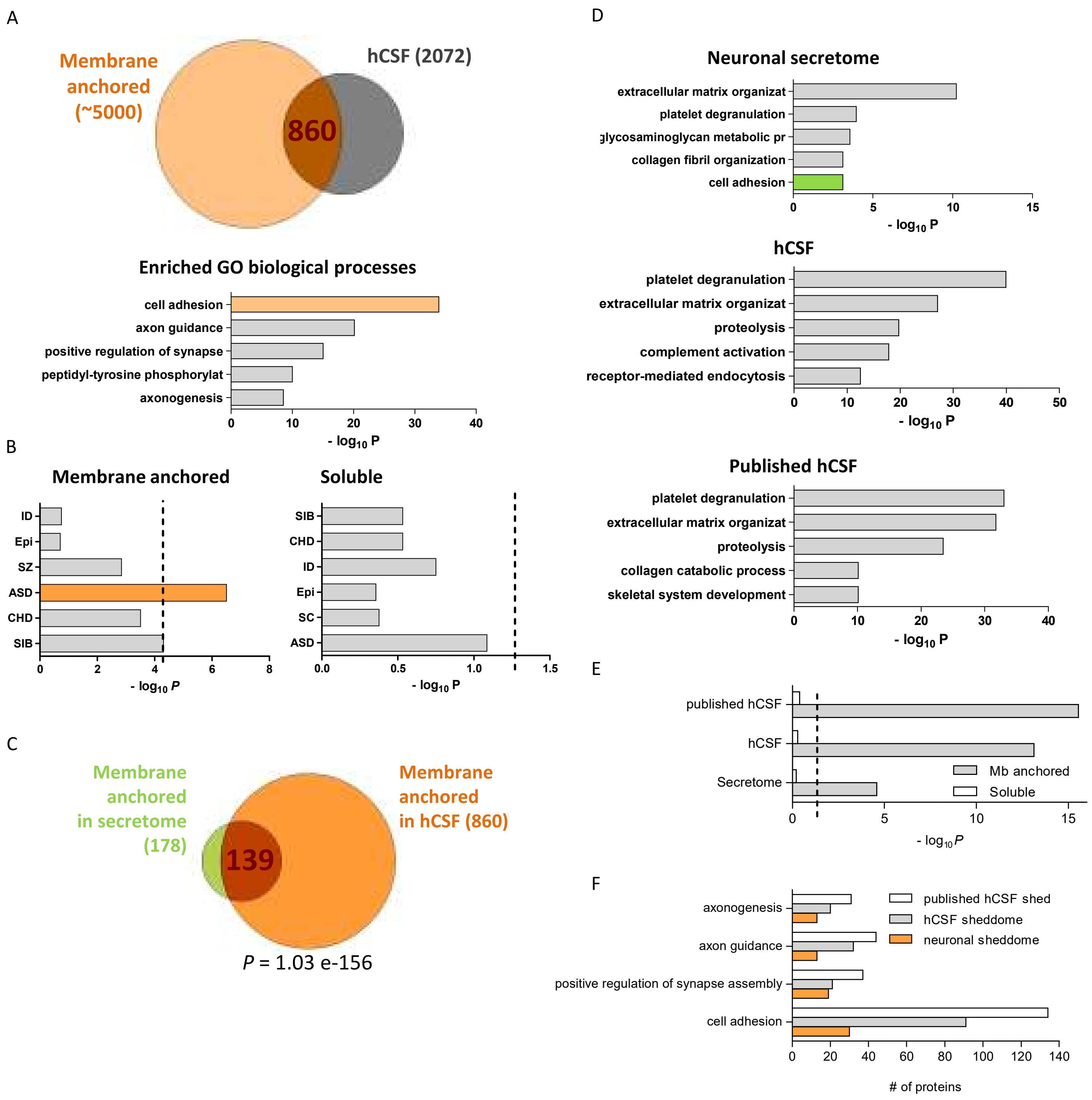

### Fig.S2

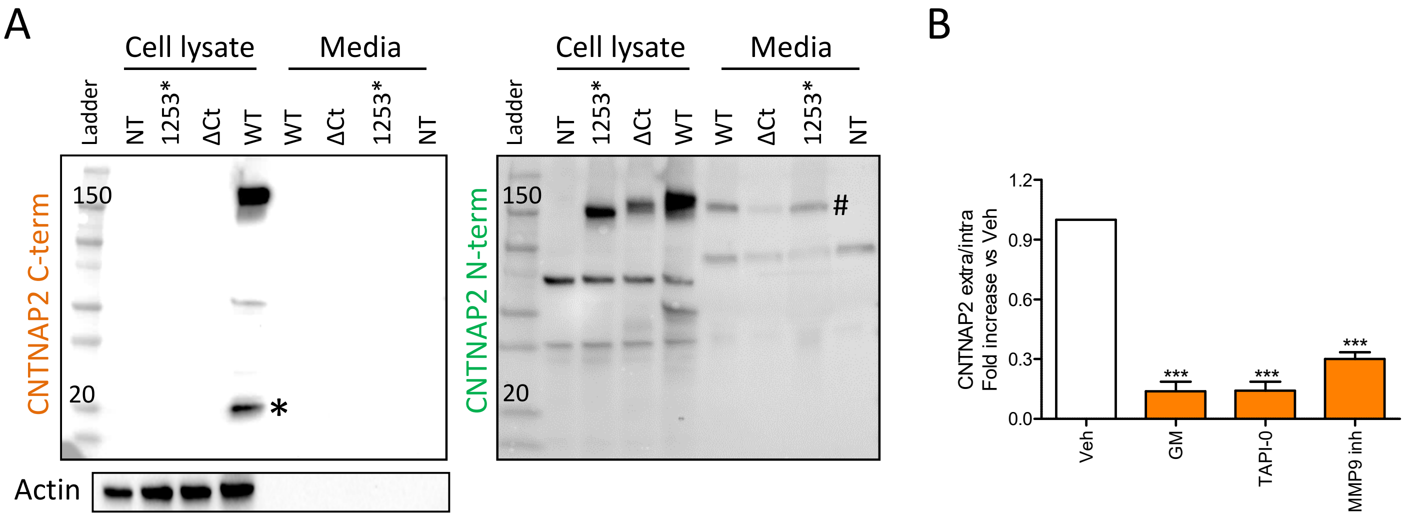

### Fig.S3

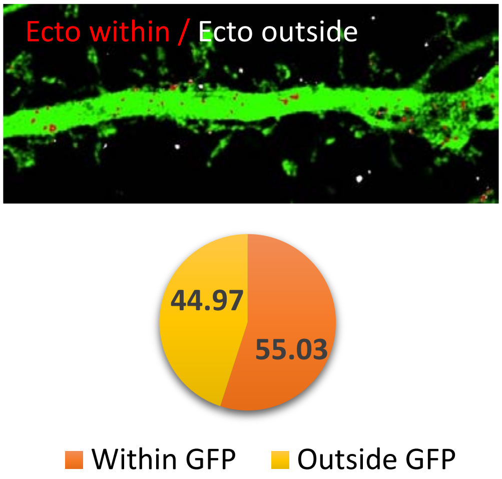

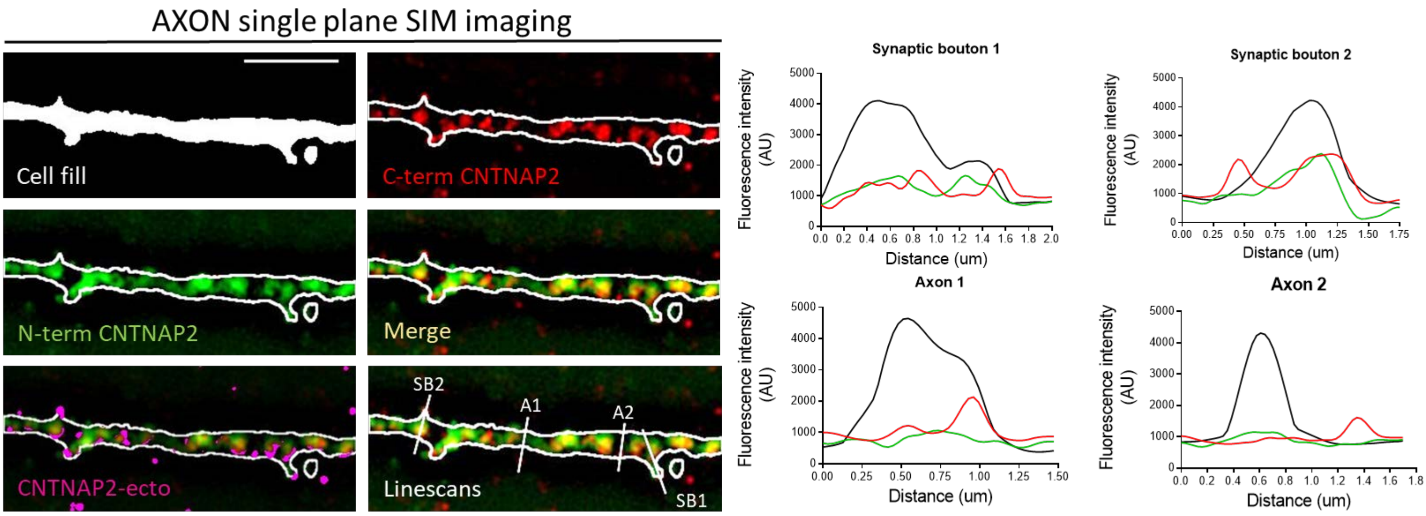

### Fig.S4

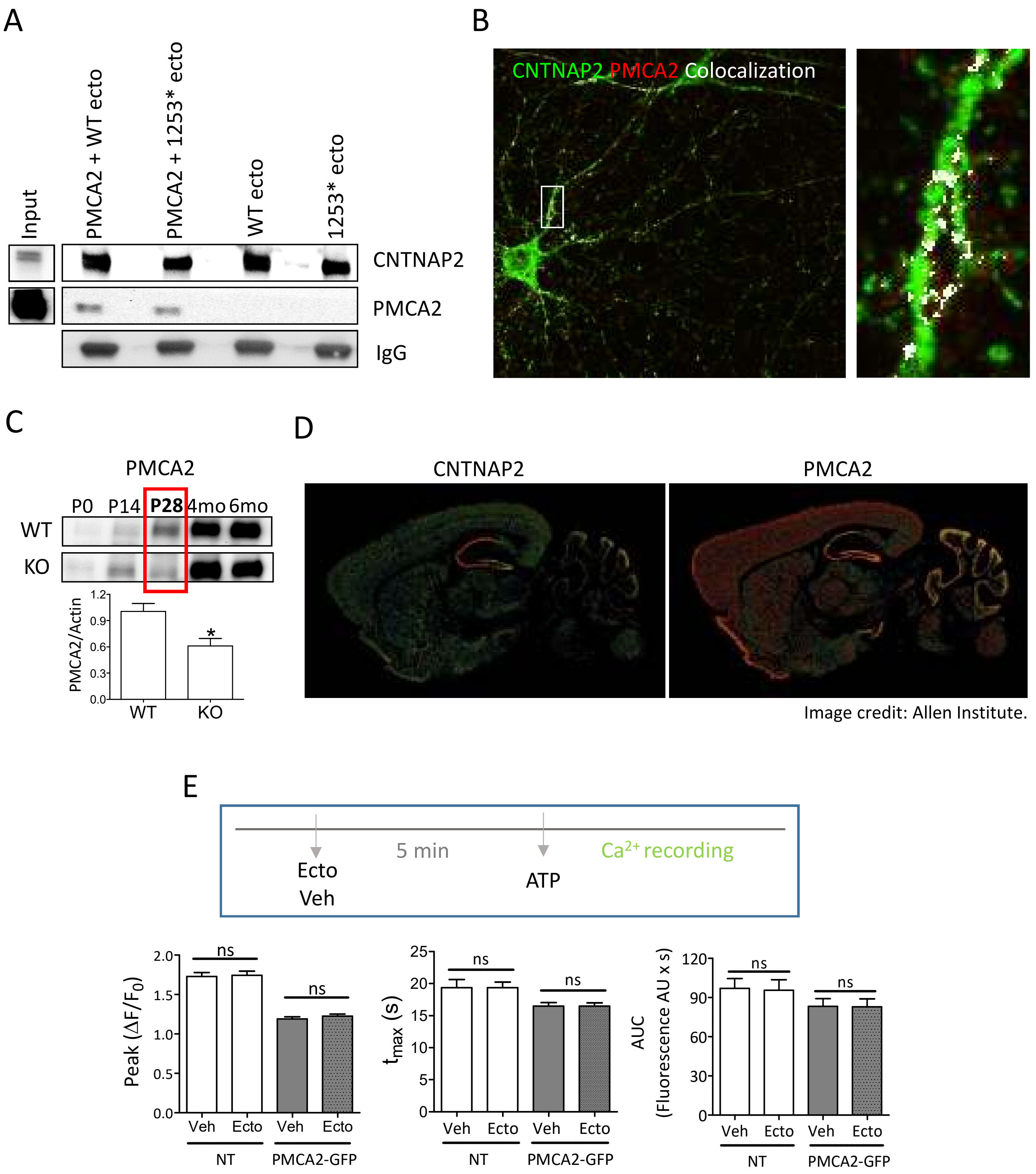

### Fig.S5

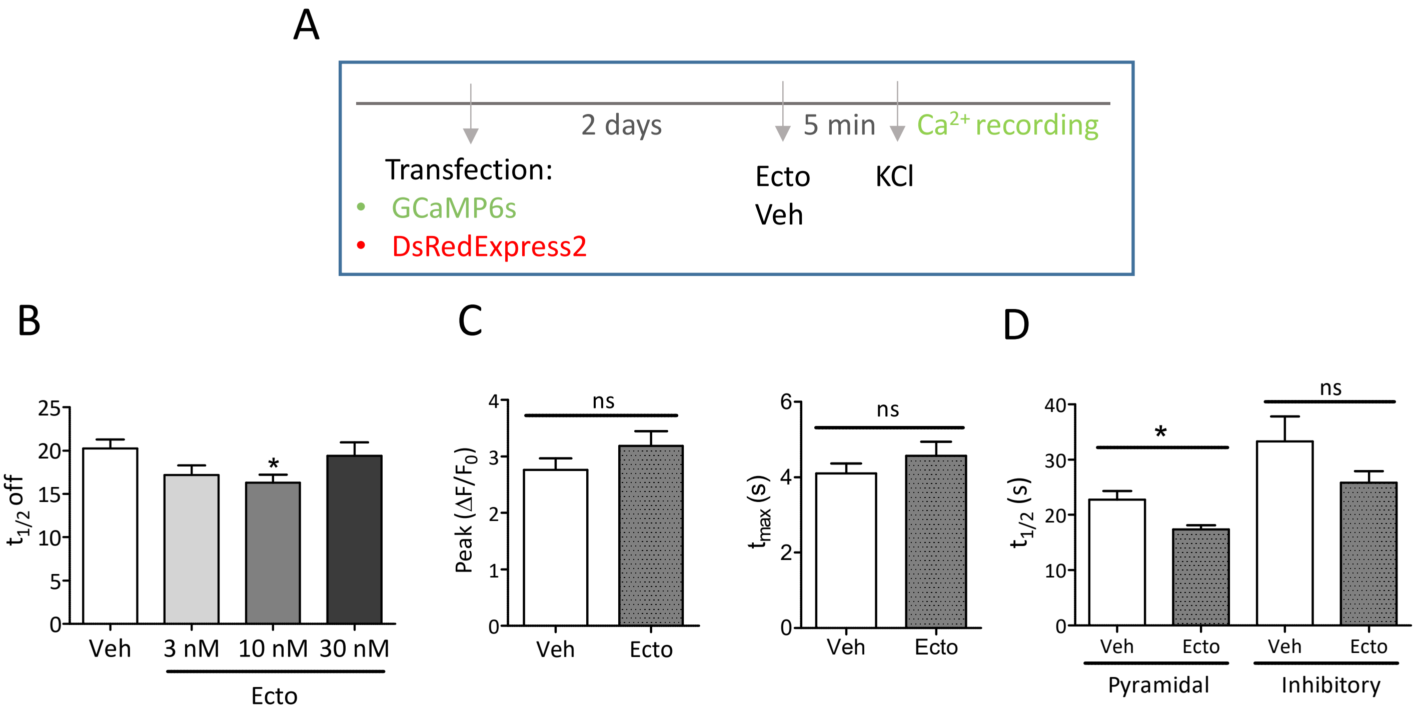

### Fig.S6

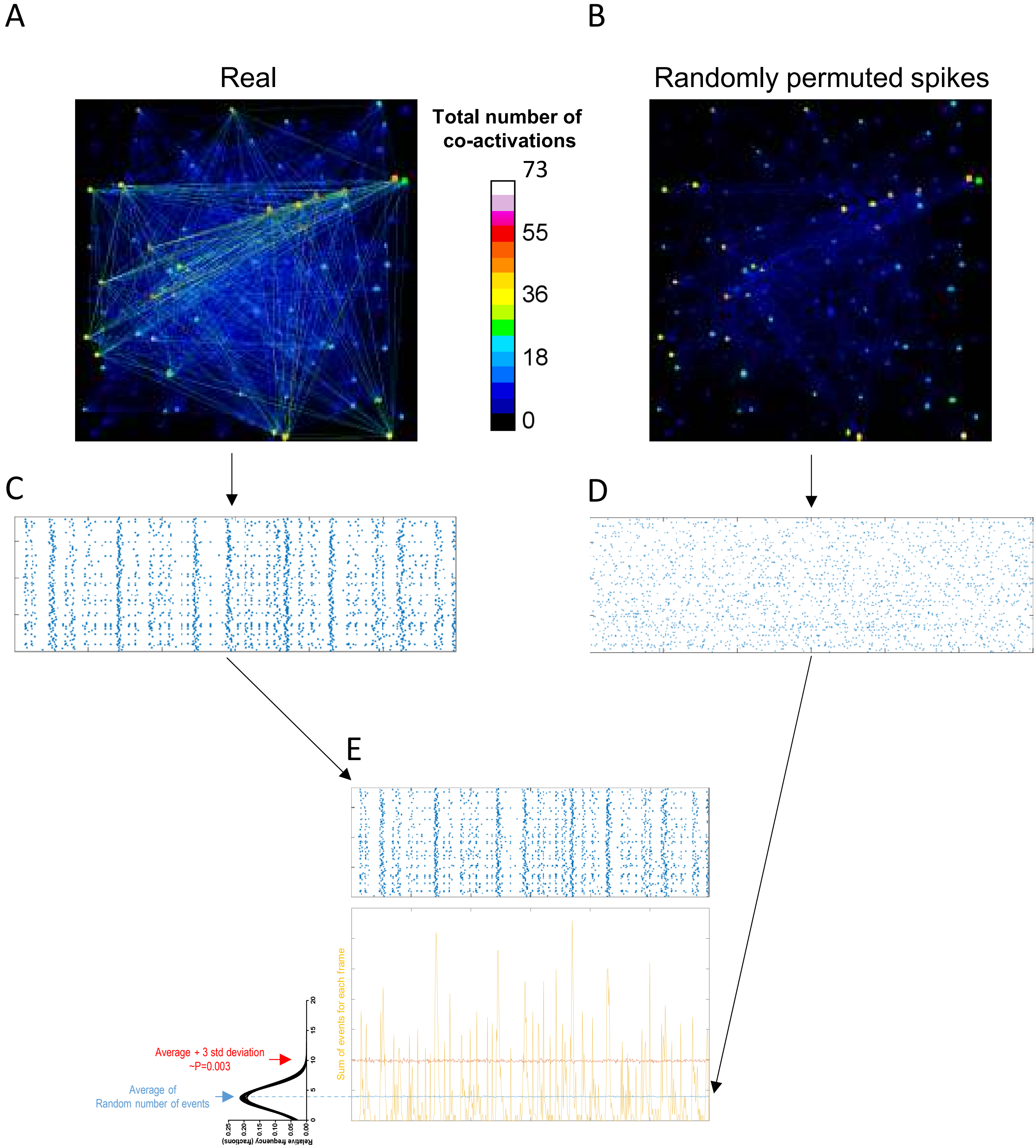
